# HiC-GNN: A Generalizable Model for 3D Chromosome Reconstruction Using Graph Convolutional Neural Networks

**DOI:** 10.1101/2021.11.29.470405

**Authors:** Van Hovenga, Oluwatosin Oluwadare, Jugal Kalita

## Abstract

Chromosome conformation capture (3C) is a method of measuring chromosome topology in terms of loci interaction. The Hi-C method is a derivative of 3C that allows for genome wide quantification of chromosome interaction. From such interaction data, it is possible to infer the three-dimensional (3D) structure of the underlying chromosome. In this paper, we use a node embedding algorithm and a graph neural network to predict the 3D coordinates of each genomic loci from the corresponding Hi-C contact data. Unlike other chromosome structure prediction methods, our method can generalize a single model across Hi-C resolutions, multiple restriction enzymes, and multiple cell populations while maintaining reconstruction accuracy. We derive these results using three separate Hi-C data sets from the GM12878, GM06990, and K562 cell lines. We also compare the reconstruction accuracy of our method to four other existing methods and show that our method yields superior performance. Our algorithm outperforms the state-of-the-art methods in the accuracy of prediction and introduces a novel method for 3D structure prediction from Hi-C data.

**Author Summary:** We developed a novel method, HiC-GNN, for predicting the three-dimensional structures of chromosomes from Hi-C data. HiC-GNN is unique from other methods for chromosome structure prediction in that it learns in an eager setting rather than a lazy setting. Thus, the models learned by HiC-GNN can be generalized to unseen data. To the authors’ knowledge, this generalizing capability is not present in any existing methods. We show that this generalization is robust to input resolution, restriction enzyme, and contact sparsity. We also show that our method outperforms existing methods using both generalized and non-generalized models. Moreover, we also show that our method is more robust to contact variance than the compared methods.

**Availability:** All our source codes and data are available at https://github.com/OluwadareLab/HiC-GNN, and is made available as a containerized application that can be run on any platform.

## Introduction

The structure of chromosomes is known to influence several genomic functions [1], [2], [3]. Thus, discovering the three-dimensional (3D) structure of chromosomes is important for understanding the functional and regulatory elements of genomes. For this reason, chromosome conformation capturing techniques such as 3C [4], 4C [5], 5C [6], and Hi-C [7], [8], [9] were developed to analyze the spatial organization of chromatin in a cell. In general, chromosome conformation capture relies on quantification of contacts between genomic loci to give insight into the structural organization of the genome. Hi-C is a chromosome conformation capture technology that allows for all-to-all quantification of intra-genomic contacts, i.e., contacts are measured between each pair of loci within the chromosome. This is accomplished via the following steps [7–9]. First, chromatins between several chromosomes are cross linked using a fixative solution. Then, the chromatin is isolated and digested by an enzyme. This results in pairs of crosslinked DNA fragments that may differ linearly but are close in physical space. These separate fragments are then re-ligated, and the crosslinks are reversed, thus resulting in templates. These templates are then amplified and interrogated, usually using polymerase chain reaction (PCR) and DNA sequencing. The resulting data describes the frequency of ligation junctions between genomic loci. These relative contact frequencies describe the proximity of the loci in 3D space. Due to its all-to-all nature, the Hi-C method allows for global insight into the spatial organization of entire genomes.

The high quantity of data that is produced with the Hi-C method has led to the development of several computational methods that aim to make inference of the 3D structure of chromosomes from their respective Hi-C data [10]. A strategy often employed by these computational methods is the distance-restraint optimization strategy [11], [12], [13], [14], [15], [16]. Usually, the distance-restraint method converts the contacts of the input Hi-C map to distances using an inverse power law [9]. These distances are typically referred to as *wish distances*. Following this conversion step, a set of *xyz* coordinates is initialized; each *xyz* coordinate corresponds to a locus in the chromosome. The model is then trained by optimizing these **xyz** coordinates so that the pairwise Euclidean distances of the predicted structure accurately recreate the wish distances of the input.

### Motivation

There are several limitations associated with traditional distance-restraint methods. Firstly, most distance-restraint methods assume independence among separate chromosomal contacts, which is likely false due to the self-attracting nature of polymers [17]. Moreover, ignoring intra-contact correlations removes a potentially valuable source of information for structure prediction. The second limitation associated with distance-restraint methods is that, to the authors’ knowledge, all current distance-restraint methods are instance-based, i.e., the results of training a model on a given Hi-C map correspond to the nothing but the chromosome associated with the input data. This means that to predict the structure of a given chromosome, one must retrain an entirely new model. This leads to intense computational requirements when using these methods to make predictions on large data sets, such as those with high resolution. Moreover, this instance-based nature associated with traditional distance-restraint methods means that these methods tend to fail when the input data is sparse as there are fewer features that can be utilized in training.

In this paper, we present a novel distance-restraint method for 3D chromosome reconstruction from cis-chromosomal HiC contacts that addresses each of these limitations associated with traditional distance-restraint methods. Our method relies on a graphical interpretation of Hi-C data. From this graphical interpretation, we use a node embedding algorithm to generate features corresponding to each chromosomal locus. These features are then utilized to train a graph convolutional neural network (GCNN) to generate predictions of the **xyz** coordinates corresponding to each chromosomal locus.

To the authors’ knowledge, **HiC-GNN** is the only chromosome structure prediction algorithm that learns in an eager setting**. Thus, HiC-GNN is the only method designed to allows for generalizable chromosome structure prediction models.** Specifically, we show that our model can generalize across three data variations:

1. **Generalization across resolutions**: a model trained on the Hi-C map of a given chromosome at one resolution can be used to accurately predict the structure of the same chromosome using a different Hi-C map resolution as the input. This allows us to train a model on low resolution data and make predictions for high-resolution data, thereby circumventing the computational expenditure associated with training a new model on high-resolution data.
2. **Generalization across restriction enzymes**: a model trained on a Hi-C map of a given chromosome utilizing some restriction enzyme in the Hi-C experiment can be used to accurately predict the structure of the same chromosome using a Hi-C map obtained with a different restriction enzyme as an input.
3. **Generalization across cell population**: a model trained on a Hi-C map of a given chromosome corresponding to some cell population can be used to accurately predict the structure of the same chromosome using Hi-C data obtained from a different cell population. This allows us to train a model on contact-sparse data (i.e., contact maps with fewer contact frequencies) and make predictions on denser contact maps.

These generalizations allow for several benefits associated with HiC-GNN that are absent in other methods. *Generalization 1* has the practical benefit of being able to train a model on low resolution data while still being able to make predictions on high resolution data, thereby avoiding the additional computational requirements associated with training a model on high resolution data. This is particularly important since the computational requirements of some methods limit their use to low resolution data. *Generalization 2* shows that our models are robust to biases introduced by choices of restriction enzymes, i.e., we can ensure that the predicted structure of a given chromosome is consistent irrespective of which restriction enzyme was used in the training data. *Generalization 3* shows that our models are robust to contact sparsity in the data.

We validate the reconstructive performance and the generalization capabilities of our method on three separate data sets from the GM12878, GM06990, and K562 cell lines and make comparisons with four other Hi-C chromosome reconstruction methods; ShRec3D [18], ShNeigh2 [19], ChromSDE [16], and LorDG [11]. We also validate the reconstructive performance of our method using orthogonal ChIA-PET data from the GM12878 cell line.

### Overview of Other Methods

We now give a brief overview of the other methods existing in the literature and describe the methods to which we compare the performance of HiC-GNN.

There currently exists many methods for 3D chromosome reconstruction. MCMC5 is a method which uses a Markov Chain Monte Carlo (MCMC) for sampling spatial coordinates from the posterior distribution generated by interaction frequency data under a Gaussian prior [20]. BACH also uses MCMC to sample spatial coordinates, except the authors assume a Poisson distributed prior [21]. PASTIS also assumes that spatial coordinates are related to contact frequencies according to a Poisson distribution; however, spatial coordinates are optimized via maximizing the likelihood of the Poisson distribution [22]. Chromosome3D is a distance restraint method which optimizes distances using distance geometry simulated annealing [15]. LorDG is a distance restraint method that uses an objective function derived from the Lorenzian function. The Lorenzian objective smooths inconsistencies in the Hi-C due to heterogeneous cell populations by rewarding the satisfaction of consistent restraints whose value is not affected by the violation of inconsistent restraints. Finally, ChromSDE is a distance restraint method that relies on semi-definite programming to optimize the predicted structures. Moreover, ChromSDE relies on a golden search algorithm to infer the relationship between interaction frequency and distance. We chose to compare our method to LorDG and ChromSDE due to their ability to outperform several other distance and contact-based algorithms, that is use the contact data directly for 3D structure reconstruction [10], [11], [16], [23]. Thus, we can consider this methods top-performers, and representative methods for distance instance-based method for chromosome 3D structure reconstruction.

We also compare our method to ShNeigh2 and ShRec3D. Both ShNeigh2 and ShRec3D are methods that consider the neighborhood structure of contact sites in Hi-C data. Like our method, these two methods rely on a graphical interpretation of Hi-C data. This is the reason why we choose to include these two methods in our method evaluation. ShRec3D considers the neighborhood structure of contact sites by utilizing a shortest path algorithm on the Hi-C data to derive distances from contacts. The structure of the chromosome is then inferred from these distances using multi-dimensional scaling. Recently proposed method, ShNeigh, incorporates neighborhood dependence by defining an affinity matrix associated with the input contact matrix defined from a Gaussian distribution. The entries of this affinity matrix are then utilized as regularization terms in the objective that is thence optimized. The authors of ShNeigh present two versions of the algorithm, ShNeigh1 and ShNeigh2. The difference between these versions is that ShNeigh1 assumes a constant relationship between interaction frequency and distance, whereas ShNeigh2 optimizes this relationship dynamically. For this reason, ShNeigh2 is slower than ShNeigh1 but usually produces better results. We compared our method with ShNeigh2.

## Materials and Methods

The crux of our method is a graphical interpretation of the input Hi-C data. Recall that a Hi-C map for a given chromosome is an *N × N* symmetric matrix whose *ij^th^* entry corresponds to the contact frequency between locus *i* and locus *j*. *N* refers to the total amount of loci observed in the Hi-C map. Our method interprets this matrix to be an adjacency matrix corresponding to an edge-weighted, un-directed graph consisting of *N* nodes. In this formulation, the *ij*^th^ entry of a given Hi-C map denotes the edge weight between node *i* and node *j*, and zero entries imply that the nodes are not connected. This graphical interpretation of the Hi-C data allows for the topology of the graph to be considered during the reconstruction.

With this graphical interpretation, we may formulate the task of predicting the structure of the chromosome as a node regression problem. Specifically, we are given a graph with unlabeled nodes corresponding to intra-chromosomal loci and edge weights corresponding to contact frequencies between these loci. Our task is to assign **xyz** coordinates to each node such that the difference between the true chromosomal structure and predicted chromosomal structure is minimized. Fig 1 gives a high-level overview of how we utilize the graphical interpretation of Hi-C data to accomplish this task.

**Fig 1.**
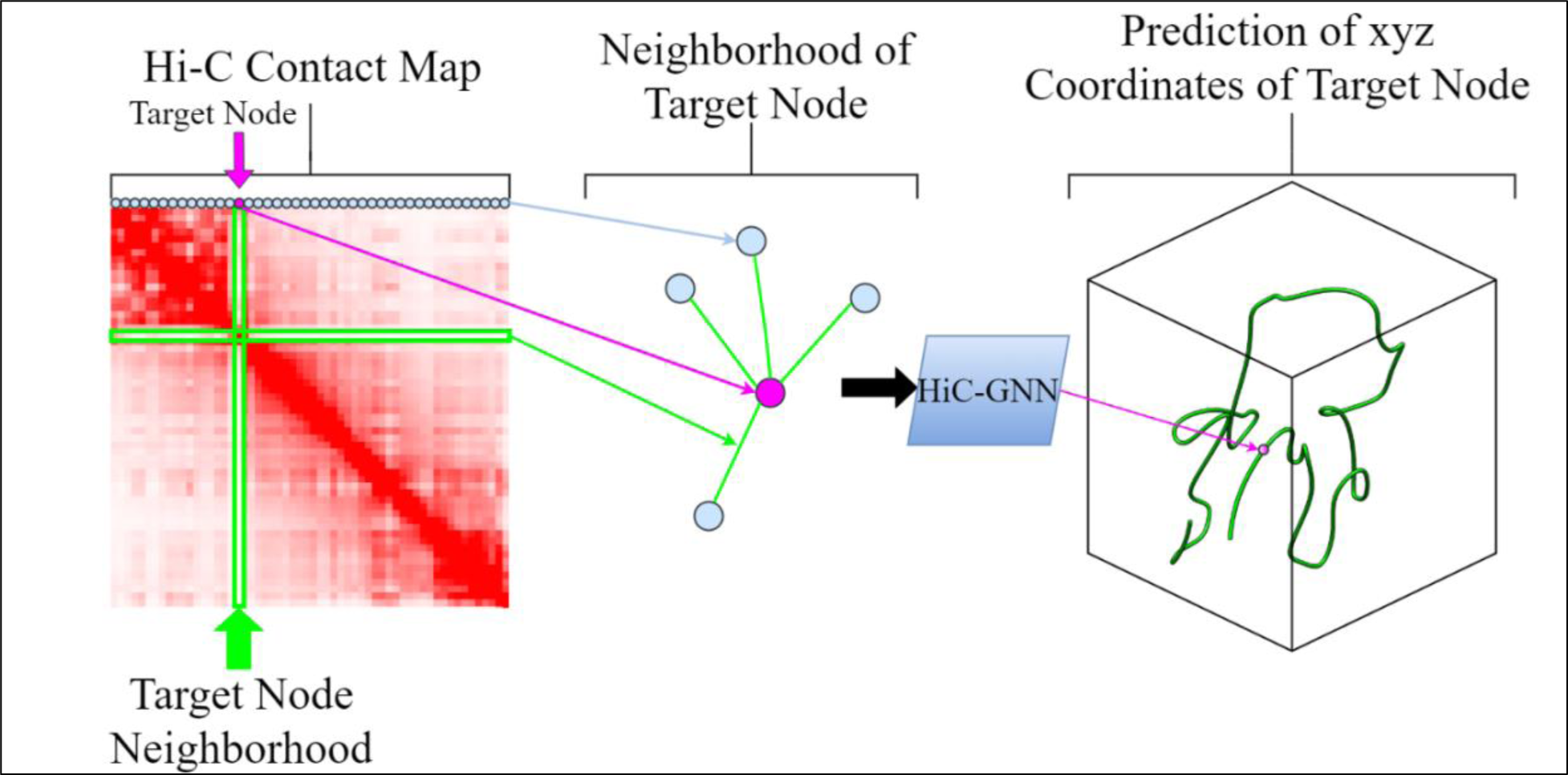
HiC-GNN 3D Chromosomal Structure Prediction Pipeline. Here we present a high-level overview of how HiC-GNN accomplishes the task of 3D chromosomal structure prediction from Hi-C data. This illustration shows takes as input a Hi-C contact map, creates a feature vector for each node and perform maximization using GCNN algorithm to generate a predicted 3D coordinate for each node depicted from the Hi-C contact map.

Our method takes a Hi-C map of cis-chromosomal contacts of a given chromosome as an input. From this map, we generate feature vectors for each node using a node embedding algorithm. We also generate ground truth, or *wish* distances, from the input map according to a standard conversion formula typical in most distance-restraint methods. We then normalize the input Hi-C map to the range [0,1] using Knight-Ruiz (KR) matrix balancing [20] to promote numerical stability in the training process. This normalization technique also mitigates biases in the Hi-C data [21]. We use this normalized map along with the node embeddings as inputs to a GCNN. The output of the GCNN is a set of *xyz* coordinates corresponding to each node of the input graph. We then compute the pairwise distances between each of these coordinates and compare to the wish distances corresponding to the input Hi-C map using mean squared error (MSE). We find the optimal coordinates by minimizing MSE through backpropagation of the GCNN. This optimization is performed using the Adam optimizer [22]. We use a convergence threshold to determine when the network is sufficiently optimized, i.e., we train until MSE is below a certain value.

### Conversion of Contacts to Wish Distances

One challenge posed by 3D chromosome structural inference is the lack of ground truth associated with the input data. We would like to optimize the output coordinates of our model to match the true pairwise distances corresponding to the loci of the input chromosome, but these true distances are generally unknown. It has been shown both empirically and theoretically, however, that relationship between the distances and contact frequencies between two loci is inversely exponential [9], [23], [24], [25]. Thus, we can estimate the true pairwise distance between locus *i* and locus *j* by using equation (1)

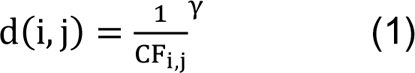

where CF_i,j_, is the interaction frequency between locus *i* and locus *j*. In general, the value for *γ* is unknown and varies depending on the underlying chromosome. It has been shown, however, that γ lies in the range [.1,2] for most, common cell types [26]. In our experiments, we assume that the optimal conversion belongs to the set {0.1, 0.2, …, 2} and perform a grid search in this interval to find the optimal model. This conversion is used in several other distance-restraint algorithms and has been shown to be a valid means for generating ground-truth distance data [27].

### Node Feature Creation

Another challenge associated with our formulation of 3D structure reconstruction as a node regression problem is the lack of features associated with the nodes we would like to regress. Hi-C data only defines a graph structure through weighted edges between featureless nodes. Thus, we must create node features to serve as inputs to the regression problem. These node features ideally have two desirable properties. Firstly, we would like these node features to be correlated to the underlying graph structure, i.e., node embeddings within regions of high connectivity should be similar. Secondly, we would like this similarity defined from the graph structure to translate to similarity of node features in Euclidean space so that the 3D structure of the chromosome can be inferred from these features. A natural way to accomplish these two goals is to create vectorized representations of each node utilizing a node embedding algorithm and use these representations as the input node features.

We create node features using the LINE node embedding algorithm [28] to be input into our GCNN. LINE is a node embedding algorithm that is specifically adapted to scalable use on large graphs. LINE has shown success in predicting chromosome compartmentalization from Hi-C data [29]. One advantage associated with LINE in the context of this specific application is that LINE considers edge weights when generating embeddings. LINE also accounts for both first and second order proximities in the input graph. Thus, the embeddings from LINE account for correlations between the contact values of the Hi-C map and preserve higher order relationships between node neighborhoods. We utilized a TensorFlow implementation of LINE available at https://github.com/shenweichen/GraphEmbedding.

### Hi-C Map Normalization

The inputs to the GCNN are a set of node features and an adjacency matrix defining node-wise connectivity. In this context, the entries of the adjacency matrix are determined by the contact frequencies in the underlying Hi-C map, the values of which are often in the hundreds of thousands. Thus, to promote numerical stability of the GCNN, we normalize the values of this adjacency matrix to the interval [0, 1]. We perform this normalization using Knight-Ruiz (KR) matrix balancing [20]. The result of Knight-Ruiz balancing is a doubly stochastic matrix. This technique has been used in several other applications of Hi-C data [21].

### Graph Convolutional Neural Network Architecture

Following the generation of node feature vectors, the regression of node *xyz* coordinates is performed using a GCNN. The advantage of utilizing a GCNN to estimate 3D coordinates from the input features as opposed to just using a standard neural network is two-fold. Firstly, GCNNs incorporate the graphical structure of the Hi-C data features, whereas standard neural networks have no way of interpreting graphical relationships from the data. Secondly, the shape of the input layer of the network depends only on the shape of the node features and is independent of the quantity of nodes in the input adjacency matrix. This independence is what allows us to generalize models between input Hi-C maps of potentially different sizes.

Our method relies on a consolidate-update inspired by the GraphSAGE algorithm [30]. In general, the consolidate-update strategy involves a consolidation of the the features of nodes in the neighborhood of a target node followed by an update of the target node’s feature via some trainable function. Assume we are computing the output of node x_i_. We first consolidate the node features of the neighborhood of x_i_ using the equation (2)

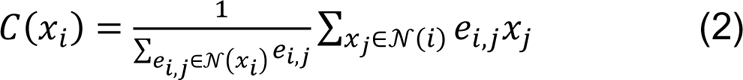

Where 𝒩(*i*) is the neighborhood of x_i_ and e_i,j_ is the edge weight between node *i* and node *j*. We then compute the updated target node **x_i_′** using equation (3)

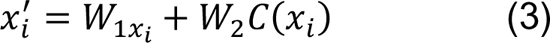

where *W*_1_ and *W*_2_ are parameter matrices that are updated utilizing backpropagation. Note that, to ensure generalizability across input maps of various node quantities, *W*_1_ and *W*_2_ are shared across all nodes. The updated node features are then passed through a four-layer multilayer perceptron (MLP) which outputs the *xyz* coordinates corresponding to the target node. Each hidden layer of the GCNN uses ReLU activation. The output layer does not use any activation so as to not restrict the domain of the predicted structure. After the GC layer, the network passes through a four-layer MLP to reduce the node features to three dimensions representing the *xyz*coordinates of each chromosomal locus. Each hidden layer of the GCNN is followed by a ReLU activation. The output layer is not followed by any activation so as to not restrict the domain of the predicted structure. We then compute the pairwise distances between each of the output *xyz* coordinates and compare these output distances to the wish distances using mean squared error (MSE). We then optimize the parameters of the network utilizing backpropagation and the Adam optimizer [22] to minimize the MSE between the distances corresponding to the output structure and the wish distances. We use a convergence threshold to determine when the network is sufficiently optimized, i.e., we train until MSE is below a certain value. The entire HiC-GNN algorithm can be visualized in Fig 2. The architecture of the GCNN can be visualized in Fig 3.

**Fig 2.**
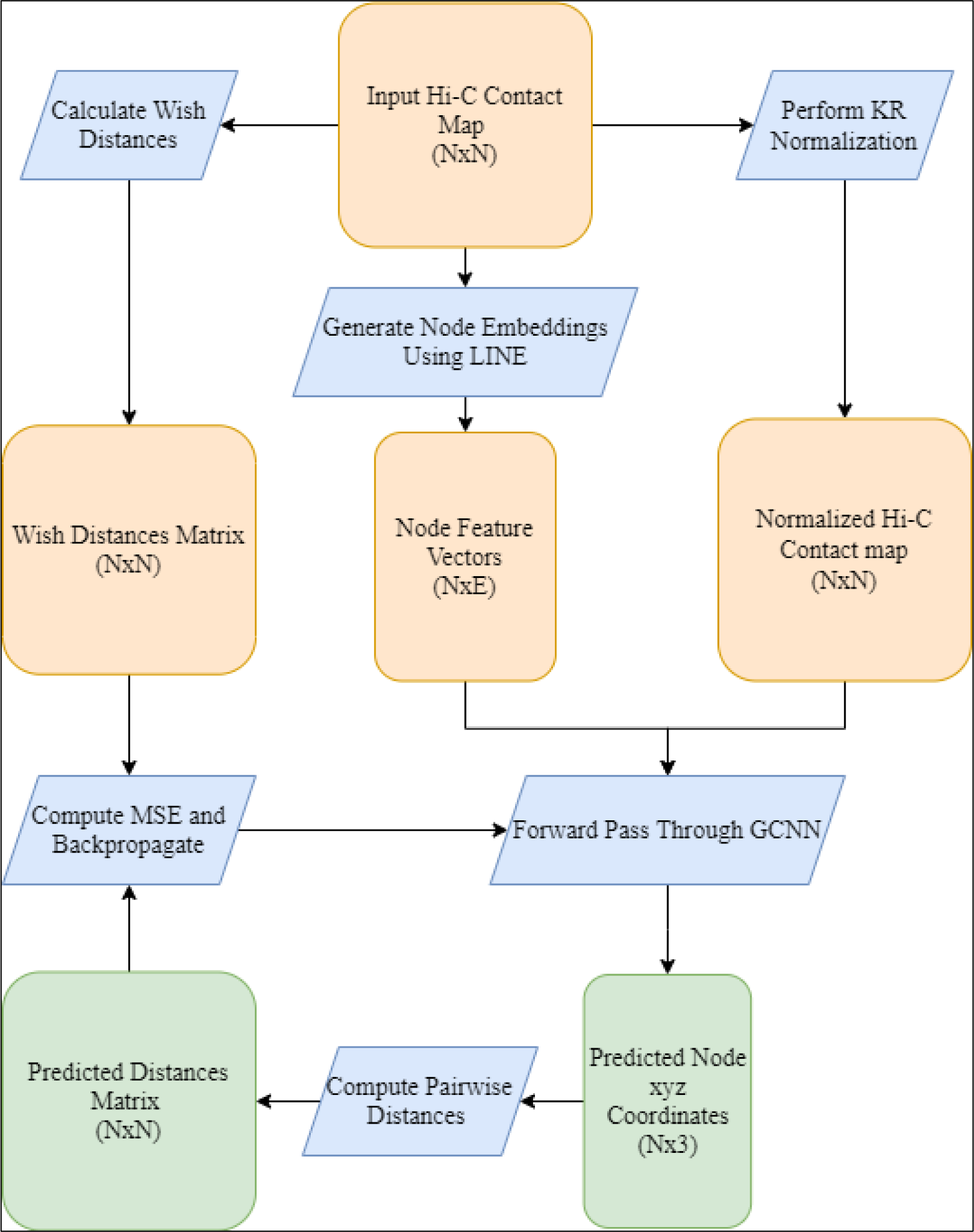
General pipeline for HiC-GNN. This figure shows the pipeline for the entire HiC-GNN algorithm. From the raw input Hi-C map, we calculate wish distances using equation (1), we generate node embeddings using the LINE algorithm, and we compute a normalized map using KR normalization. The node feature vectors and the normalized Hi-C map are then used as inputs to the graph neural network. The graph neural network is optimized by minimizing the MSE of the pairwise distances of the output structure to the wish distances. Here, *N* refers to the number of loci and *E* refers to the size of the embeddings.

**Fig 3.**
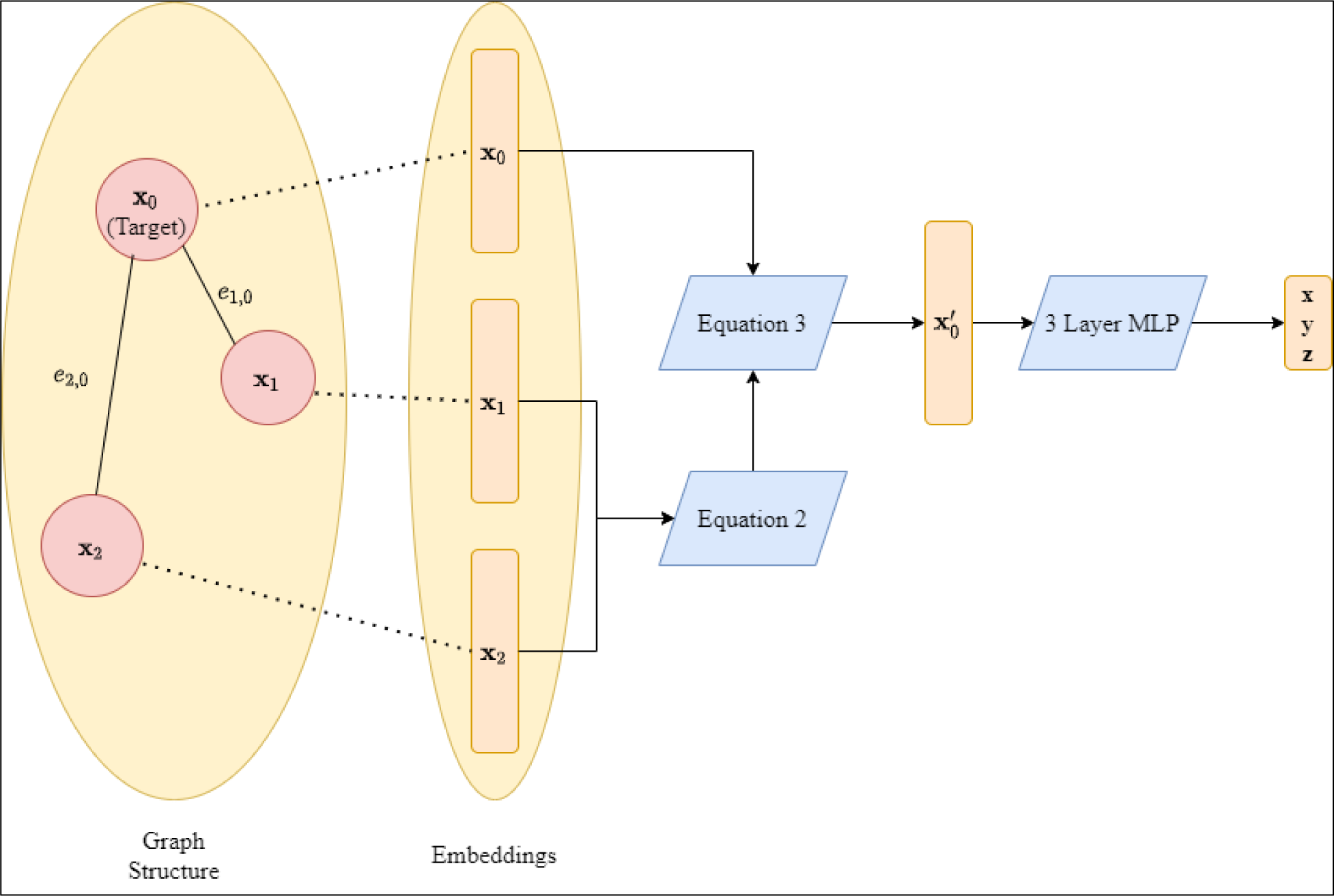
Architecture of the GCNN. This figure details the architecture of the GCNN. The node features of the target node’s neighborhood are consolidated using equation (2). The representation of the target node is then updated using equation 3. Finally, the coordinates of the target node are predicted using a 3-layer MLP. Here, we are predicting the *xyz* coordinate of **x_0_**, where *x*_1_ and *x*_2_ are the neighbors of ***x*_0_** with edge weights of *e*_1,0_ and *e*_2,0_ respectively.

### Embedding Alignment for Generalization

The process of generalizing the results of HiC-GNN involves training the GCNN on a Hi-C map and its corresponding embeddings from one set of data and utilizing this trained network to generate structures using the embeddings and maps of another set of data. It is possible, however, that the embedding distributions vary significantly across different data, thereby making generalization difficult. Thus, we assume that embeddings are only approximately similar up to isometry, i.e., we assume that the embeddings between two separate chromosomes are approximately equivalent up to rotation, translation, and scaling irrespective of the restriction enzyme, cell population, and resolution of the maps. To test this assumption, we employ an embedding realignment procedure prior to testing a generalized model on new embeddings.

Assume we have two *N* × *E* embeddings matrices, *A* and *B*. Here, *N* refers to the number of chromosomal loci and *E* refers to the embedding size. We would like to find a linear transformation that minimizes the Euclidean distance between *A* and *B*. Formally, we would like to compute (4).

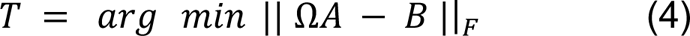

|| · ||_F_ denotes the Frobenius norm. If we impose the constraint that the matrix *T* is orthogonal, our problem is equivalent to the Orthogonal Procrustes Problem (OPP) [30]. Computing the matrix *T* in the OPP is equivalent to computing the singular value decomposition of the matrix *M* = *BA*^*T*^ [31]. Thus, the task of embedding realignment has a closed form solution and requires no additional training. Note that, in our applications, it is not guaranteed that the embedding matrices *A* and *B* have the same size due to differing numbers of chromosomal loci across differing resolutions. For this reason, we employ a simple binning procedure to match the number of rows in the embedding matrices which we describe below.

### Binning Procedure for Feature Alignment

The alignment procedure used in our model generalization assumes the existence of a linear transformation between the embedding spaces of two distinct Hi-C maps. In the case of generalizing across resolutions, however, we run into the issue of the dimensions of these spaces differing. Specifically, if *A* denotes the embedding matrix corresponding to the lower resolution data and *B* denotes the embedding matrix corresponding to the higher resolution data, then Ω*A* − *B* is not well defined since the number of rows in *A* is less than the number of rows in *B*. We fix this problem using the following binning procedure.

Assume we are performing a resolution generalization of a given chromosome. In our experiments, *A* always corresponds to the map at 1mb resolution and *B* either corresponds to the map at 500kb or 250kb resolution. This implies that *B* either as twice or four times the number of rows of *A*. See Table 1 for a visual representation of why this is the case.

**Table 1.**
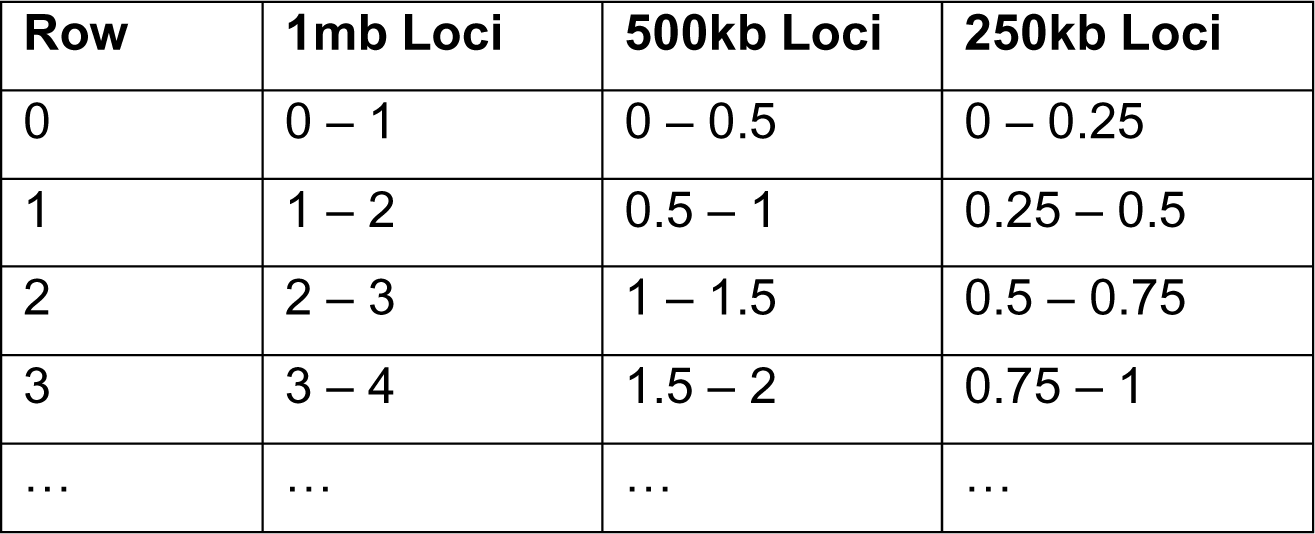
Table showing which rows of the embedding matrices correspond to which interaction sites.

The row column represents the row of an arbitrary embedding matrix. The loci columns depict which interaction sites the row of the embedding matrix corresponds to at a given input resolution. These values are given in millions of base pairs. For example, embedding of the first row of an embedding matrix generated from 1mb data corresponds to the portion of the chromosome between base pair 0 and base pair 1,000,000. The same row corresponds to the portion of the chromosome between base pair 0 and base pair 500,000 for an embedding matrix generated from 500kb data, and base pair 0 and base pair 250,000 for an embedding matrix generated from 250kb data. To force the embeddings matrices to have the same number of rows, we simply concatenate additional rows of the 1mb data such that the chromosomal region of the equivalent rows in the higher resolution embeddings matrix is contained in the chromosomal region of the given row in the 1mb embeddings matrix. See Tables 2 and 3 for an example of this binning procedure applied to the 500kb case and the 250kb case. By expanding the 1mb embeddings matrix in this way, we ensure that the dimensions of matrix *A* and matrix *B* match in the allignment procedure. Moreover, we ensure that the corresponding rows between these two matrices come from the same regions in the chromosome.

**Table 2.** Table showing how we expand the 1mb embeddings matrix to match the shape of the 500kb embeddings matrix.

| Row | Expanded 1mb Loci | 500kb Loci |
| --- | --- | --- |
| 0 | 0 – 1 | 0 – 0.5 |
| 1 | 0 – 1 | 0.5 – 1 |
| 2 | 1 – 2 | 1 – 1.5 |
| 3 | 1 – 2 | 1.5 – 2 |
| ... | ... | ... |

**Table 3.** Table showing how we expand the 1mb embeddings matrix to match the shape of the 250kb embeddings matrix. Note that the alignment process for Hi-C data often yields regions with no contacts. For sake of reducing the size of these data, many Hi-C maps simply do not include these contacts. In order to circumvent this issue, we include zero contacts in this binning procedure so that it is guaranteed the number of loci for higher resolution is a scalar multiple of the number of loci for the lower resolution.

| Row | Expanded 1mb Loci | 500kb Loci |
| --- | --- | --- |
| 0 | 0 – 1 | 0 – 0.25 |
| 1 | 0 – 1 | 0.25 – 0.5 |
| 2 | 0 – 1 | 0.5 – 0.75 |
| 3 | 0 – 1 | 0.75 – 1 |
| ... | ... | ... |

### Hyperparameter Optimization

Prior to generating results on real Hi-C data, we tuned the hyperparameters of HiC-GNN by performing a grid search on the simulated Hi-C data from Trussart et al [27]. The Trussart et al. dataset consists of multiple Hi-C maps generated from the simulation of the Hi-C protocol on multiple worm-like chain (WLC) chromosome models at varying levels of noise and structural variability. The advantage of using simulated data for hyper-parameter tuning is that unlike in the case of real Hi-C data, the structure of the chromosome is known, thereby allowing us to make a direct comparison between the outputs of HiC-GNN and the true distances of the chromosome. By optimizing the hyper-parameters of our model in a setting in which the outputs can be compared with a known structure, we ensure that our model will perform well on data where the true structure of the input chromosomes is unknown as well.

We performed our experiments on a simulated chromosome of minimal structural variability with a corresponding simulated Hi-C map involving zero noise. We chose this particular chromosome-noise configuration so that the optimal parameters selected by the grid search were not influenced by randomness associated with high levels of structural variability or noise. In our grid search, we aimed to optimize the node embedding size, the sizes of the hidden layers of the GCNN, the learning rate, and the convergence threshold. The results of this grid search can be found in tables 4 and 5. The optimal parameters are shown in bold. A spreadsheet containing all of the dSCC values for the different configurations for hyperparameter tuning can be found in the supplementary file S1 Table.

**Table 4.**
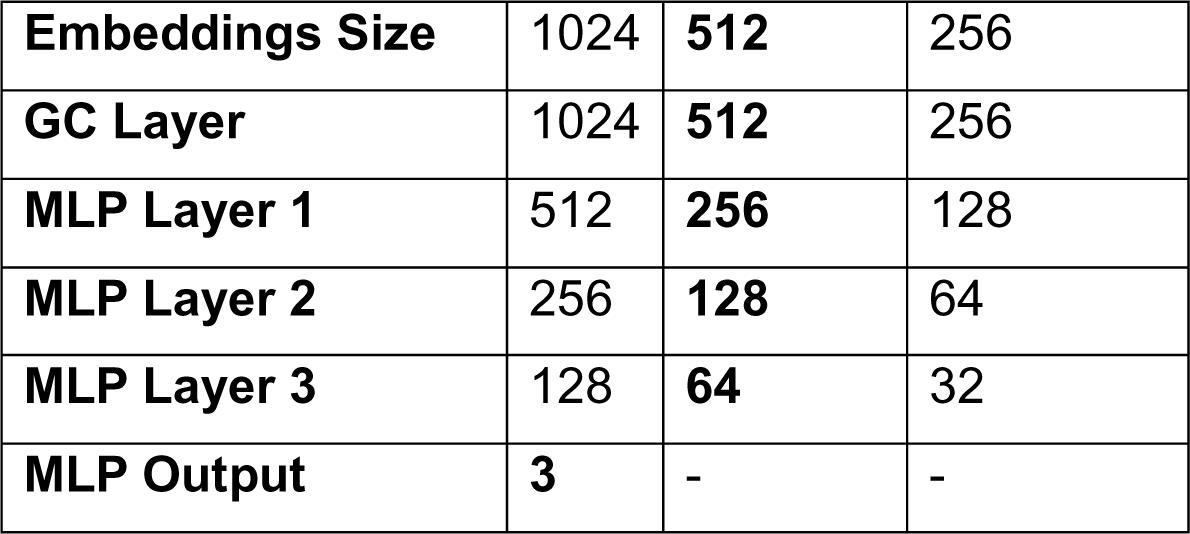
Optimal layer sizes as determined by the grid search on the simulated data. Note that the MLP must have an output of size 3 to correspond to the xyz coordinates of the chromosomal loci. The selected network settings based on the grid search are in bold on the table.

|  |  |  |  |
| --- | --- | --- | --- |
| **Embeddings Size** | 1024 | **512** | 256 |
| **GC Layer** | 1024 | **512** | 256 |
| **MLP Layer 1** | 512 | **256** | 128 |
| **MLP Layer 2** | 256 | **128** | 64 |
| **MLP Layer 3** | 128 | **64** | 32 |
| **MLP Output** | **3** | - | - |

**Table 5.** Optimal learning rate and convergence threshold as determined by the grid search on the simulated data. We explored different learning rate and convergence thresholds; the selected network settings based on the grid search are in bold on the table.

|  |  |  |  |  |
| --- | --- | --- | --- | --- |
| **Learning Rate** | 0.1 | 0.01 | **0.001** | 0.0001 |
| **Convergence Threshold** | $10^{-2}$ | $10^{-4}$ | **$10^{-5}$** | $10^{-12}$ |

## Results

### Data

#### Real Hi-C Data

To test the reconstructive performance of HiC-GNN, we utilized three data sets consisting of real Hi-C data. The first data set corresponds to the human GM12878 cell line from Rao et al. [32]. This data set consists of the Hi-C maps of 23 chromosomes generated from the Mbol restriction enzyme at 1mb, 500kb, and 250kb resolutions. This data set was downloaded from the Genome Structure Database (GSDB) repository [33] under the GSDB ID OO7429SF. We utilized this data set to test Generalization 1. The second data set corresponds to the human GM06990 cell line from Lieberman et al. [9]. This data set consists of the Hi-C maps of 22 chromosomes generated from the Ncol and HindIII restriction enzymes at 1mb resolution. We utilized this data set to test Generalization 2. The third data set corresponds to the human K562 cell line from Rao et al. [32]. This data set consists of several Hi-C maps of 23 chromosomes generated from the Mbol restriction enzyme at 1mb resolution. The genome-wide maps of this data set vary in their total number of contacts, ranging from 53 million to 932 million. This data set was downloaded from the Juicebox tool developed by Durand et al. [34]. We utilized this data set to test generalization 3.

#### ChIA-PET Data

Chromatin immunoprecipitation (ChIP) is a technique to investigate protein specific interactions in chromosomes. ChIP relies on antibodies to precipitate specific proteins, histones, or transcription factors from cell populations. ChIP can also be combined with sequencing technologies to quantify these interactions [35]. Chromosome Interaction Analysis by Paired-End Sequencing (ChIA-PET) [36] is an example of such a technology. The main difference between ChiA-PET and Hi-C data is that the ChiA-PET technique measures interactions of a unique protein in the chromosome, whereas the Hi-C technique measures interactions between any proteins in the chromosome.

To further validate our results on the real Hi-C data, we compare the outputs of our method when using Hi-C data to the interaction frequencies of an orthogonal ChiA-PET data set. We performed this validation using ChIA-PET data from the NCBI GEO database (GEO accession: GSE72816) for the RNAPII ChIA-PET data from human GM12878 cells [37]. This data measures interactions between the RNA polymerase II multicomplex; a protein complex that is responsible for gene transcription.

### Evaluation

To validate the reconstructive accuracy of our method, we use Spearman Correlation Coefficient (dSCC). dSCC is a non-parametric measure of rank correlation. The advantage to evaluating reconstructive performance using a ranked measure of similarity is that the measure is both scale and shift invariant with respect to the *xyz* coordinates of the output structure. Moreover, Trussart et al. showed that the dSCC between output structures and wish distances is a good proxy for model accuracy of distance-restraint chromosome reconstruction methods [27]. In general, dSCC values closer to 1 imply higher reconstructive accuracy. The formula for dSCC is given by equation (5).

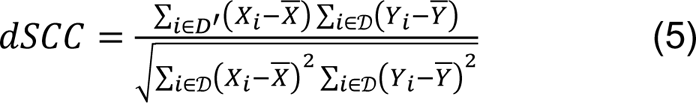

𝒟^′^ is the set of pairwise distances between all loci of the generated model, *X*_*i*_ is the rank of distance *i* in 𝒟^′^, 𝒟 is the set of wish distances corresponding to the input contact frequencies of the chromosome, and *Y*_*i*_ is the rank of wish distance *i* in 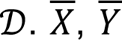 are the mean of their corresponding ranked vectors in 𝒟^′^ and 𝒟 respectively.

Note that dSCC is a non-parametric measure of rank correlation. The advantage to evaluating reconstructive performance using a ranked measure of similarity is that the measure is both scale and shift invariant. Intuitively, the model may output a perfect match of the chromosome, but the *xyz* coordinates may be scaled by a constant and rotated in space. This scaling and rotation would be accounted for in a non-ranked measure of correlation and would likely decrease the correlation value. This decrease in correlation would falsely imply that the generated model is inaccurate when the only dissimilarity between it and the ground truth is the scale and location in space. Since the purpose of modeling the chromosome in 3D space is solely for visualization, the scale and orientation of the output should not matter. Thus, dSCC is an appropriate measure of structural similarity in this context. Moreover, the simulated dataset from Trussart et al. showed that the dSCC between output structures and wish distances is a good proxy for model accuracy of distance-restraint chromosome reconstruction methods [27].

### GM12878

#### Generalization 1: Generalization Across Input Resolution

To test the reconstructive performance of HiC-GNN on real data, we evaluated the distance Spearman Correlation Coefficient (dSCC) of outputs when evaluated on Hi-C maps from the GM12878 cell line generated with Mbol restriction enzyme. To test for the effects of variability in resolution, we generated models on three separate resolutions: 1mb, 500kb, and 250kb. We compared the dSCC of our output models to the dSCC of the output models of the four other methods using the optimal hyper-parameters suggested by the authors of both methods.

We also utilized the GM12878 cell line to test how well HiC-GNN can generalize across input resolutions. To do this, we generated embeddings for one chromosome at 1mb, 500kb, and 250kb resolutions. We then trained our GCNN using the contact maps and corresponding embeddings of the 1mb data until convergence is met and stored the optimal conversion factor. Following this training, we aligned the embeddings of the 500kb and 250kb data to those of the 1mb data. We then generated structures using these aligned embeddings and their corresponding Hi-C contact maps as inputs to the pre-trained GCNN. Finally, we calculated the dSCC between the pairwise distances of the generated structures and the wish distances calculated from the input contact maps using the optimal conversion found in the training on the 1mb data.

Figs 4 and 5 show a comparison between the output dSCC values of the generalized HiC-GNN models and the output dSCC values of the non-generalized HiC-GNN models on the 500kb and 250kb data respectively. By generalized models, we mean models that were trained on the 1mb data and tested on the higher resolutions data. By non-generalized models, we mean models that were trained and tested on data of the same resolution. In these figures, we also include the output dSCC of the generalized HiC-GNN models using un-aligned embeddings as inputs to show the effect of the alignment procedure on reconstructive performance. From these figures, two things are clear. Firstly, the embedding alignment procedure increases the reconstructive performance of HiC-GNN. This suggests that the assumption of approximate similarity up to isometry of node embeddings is valid. Secondly, although there is some decrease in dSCC associated with the generalized models, most of the values are above .8 for the 500kb generalization and above .7 for the 250kb generalization. This suggests that HiC-GNN is indeed generalizing to these higher resolution data.

**Fig 4.**
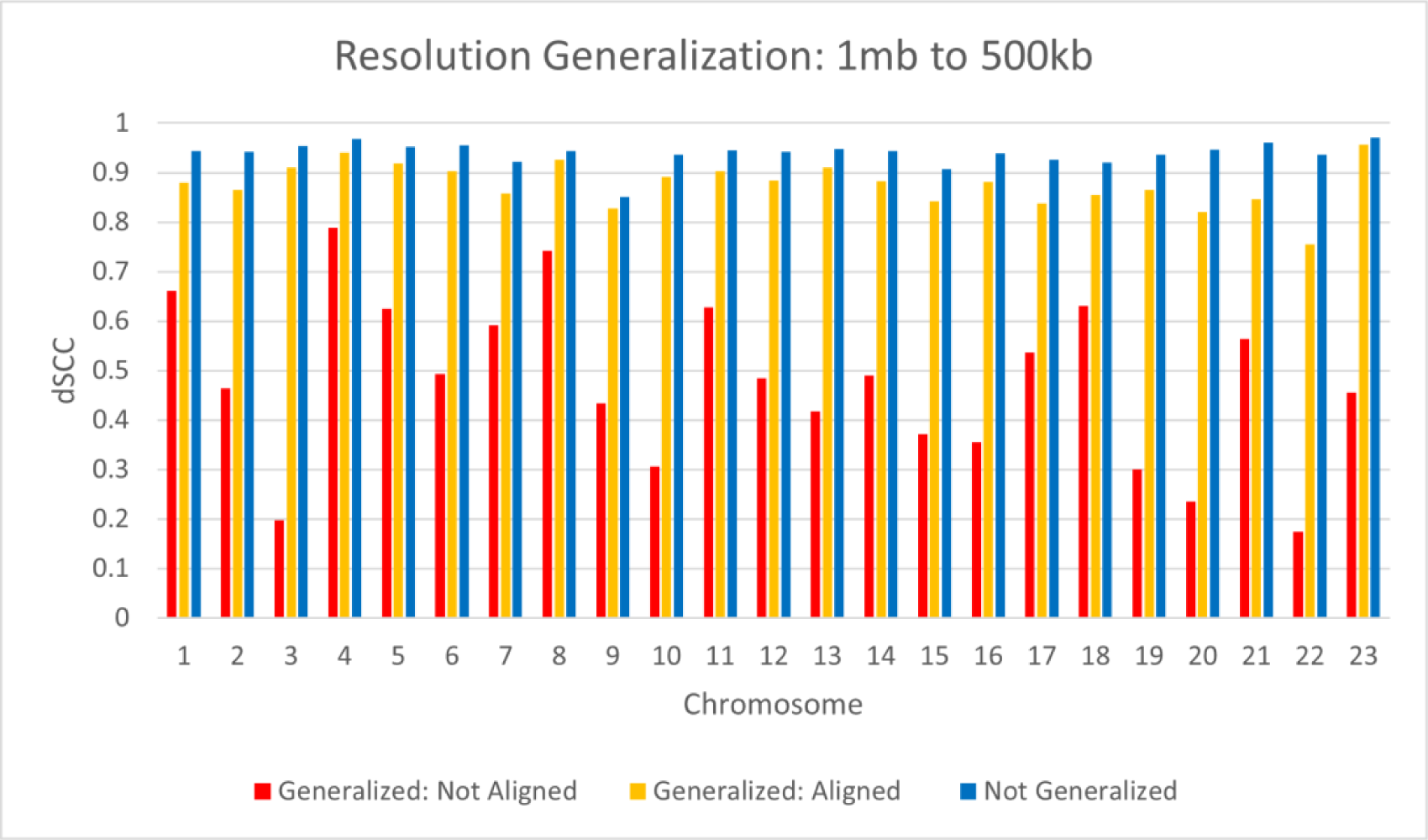
dSCC Comparison: Generalized and Non-Generalized Models at 500kb Resolution. The figure shows the dSCC values for generalized and non-generalized HiC-GNN models at 500kb resolution both with and without aligned node embeddings. The generalized models were trained on 1mb data. The difference in dSCC values between the aligned and non-aligned embeddings implies that the alignment procedure has a positive effect on reconstructive performance. The high dSCC values of the generalized models also imply that the HiC-GNN models can generalize to higher resolution data.

**Fig 5.**
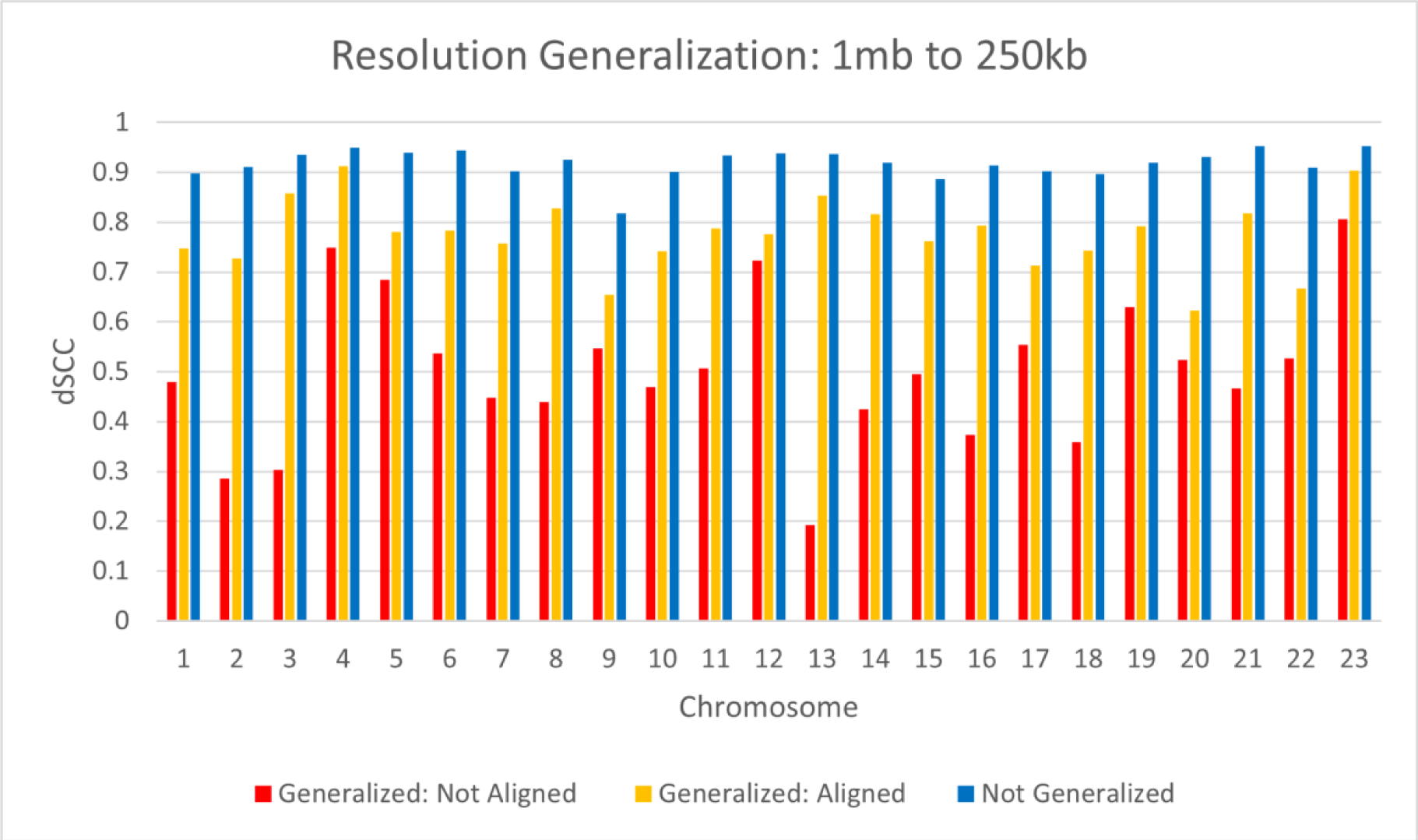
dSCC Comparison: Generalized and Non-Generalized Models at 250kb Resolution. The figure shows the dSCC values for generalized and non-generalized HiC-GNN models at 250kb resolution both with and without aligned node embeddings. The generalized models were trained on 1mb data. The difference in dSCC values between the aligned and non-aligned embeddings implies that the alignment procedure has a positive effect on reconstructive performance. The high dSCC values of the generalized models also imply that the HiC-GNN models can generalize to higher resolution data.

Fig 6 shows the comparison of HiC-GNN with the four other methods on 1mb data. Figs 7 and 8 show this same comparison along with the results from the generalized HiC-GNN models. Note that there are several missing data points for ChromSDE on the 500kb and 250kb data due to computational restraints associated with running the algorithm on these larger data sets. From these figures, it is clear that the non-generalized HiC-GNN either outperforms or is on par with the other methods. Moreover, the generalized HiC-GNN models either outperform or are on par with ShRec3D and ShNeigh. This is significant because the generalized HiC-GNN models were trained on half as many data instances in the case of the 500kb generalization and one quarter as many data instances in the case of the 250kb generalization. It is also worth noting that the dSCC values of HiC-GNN have less variation than most of the other methods. One of the main causes of variation in the dSCC values is high variance in the contact data. Fig 9 shows the variance in the contacts for each chromosome in this data set. Note that the dSCC values of chromosomes with high contact variance are significantly reduced for most of the other methods, particularly on chromosome 22. This shows that HiC-GNN is also more robust to variance in the underlying contact data. Figs 10 and 11 show the output structures of HiC-GNN corresponding to the generalized models and the original models for chromosomes 3, 11, and 13 and 2, 3, and 14 respectively. One can see that the generalized structures are indeed qualitatively similar to the non-generalized structures.

**Fig 6.**
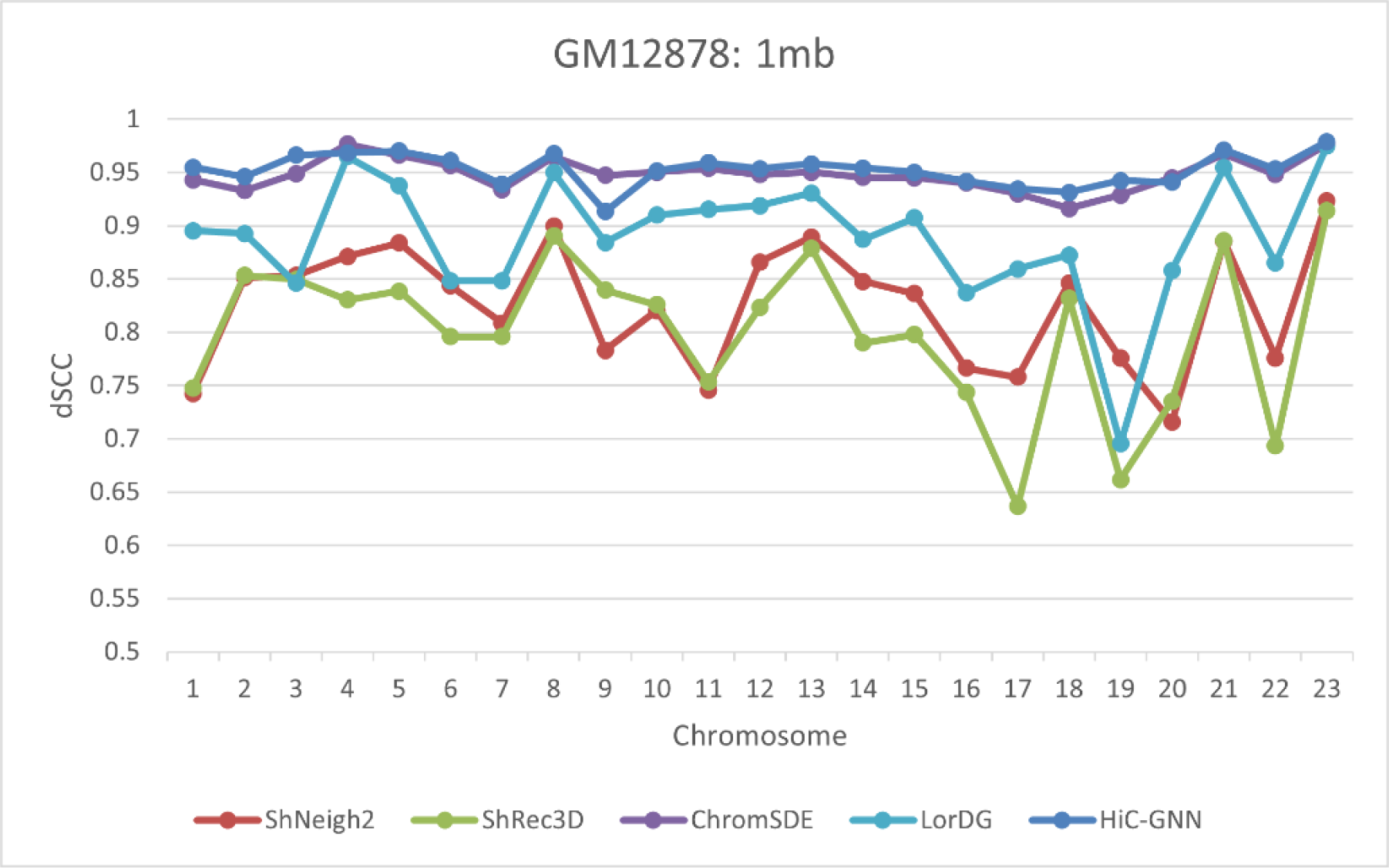
dSCC Comparison: 1mb Resolution. The figure shows a comparison of HiC-GNN with the other methods on the 1mb GM12878 cell data. HiC-GNN is either on-par or outperforms the other methods on the majority of the chromosomes.

**Fig 7.**
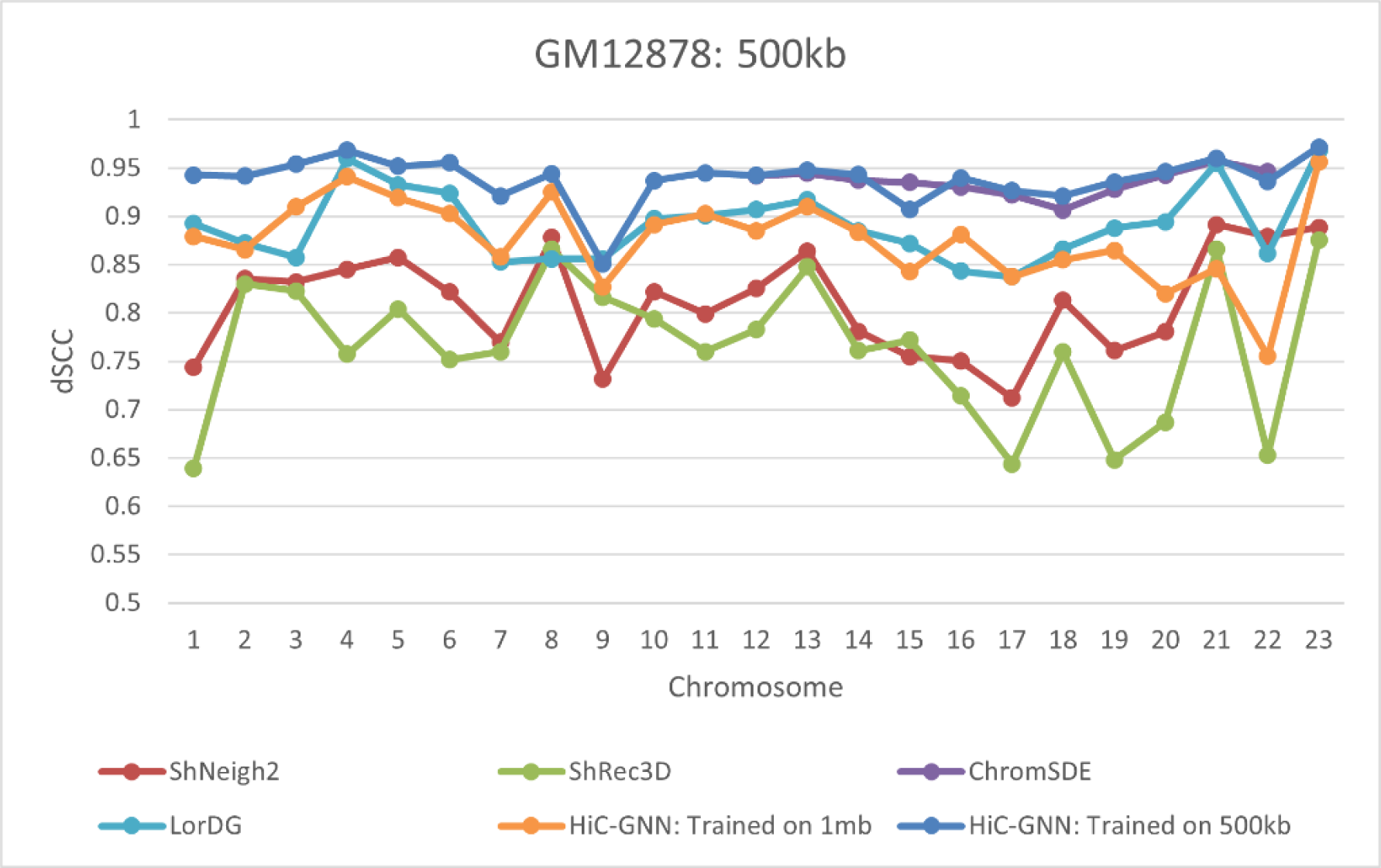
dSCC Comparison: 500kb Resolution. The figure shows a comparison of HiC-GNN with the other methods on the 500kb GM12878 cell data. The HiC-GNN models that were trained on the 500kb data are either on-par or outperform the other methods on the majority of the chromosomes. Although the HiC-GNN models that were trained on the 1mb data have lower performance, they still outperform ShNeigh and ShRec3D and are on par with LorDG despite being trained on half as many data instances.

**Fig 8.**
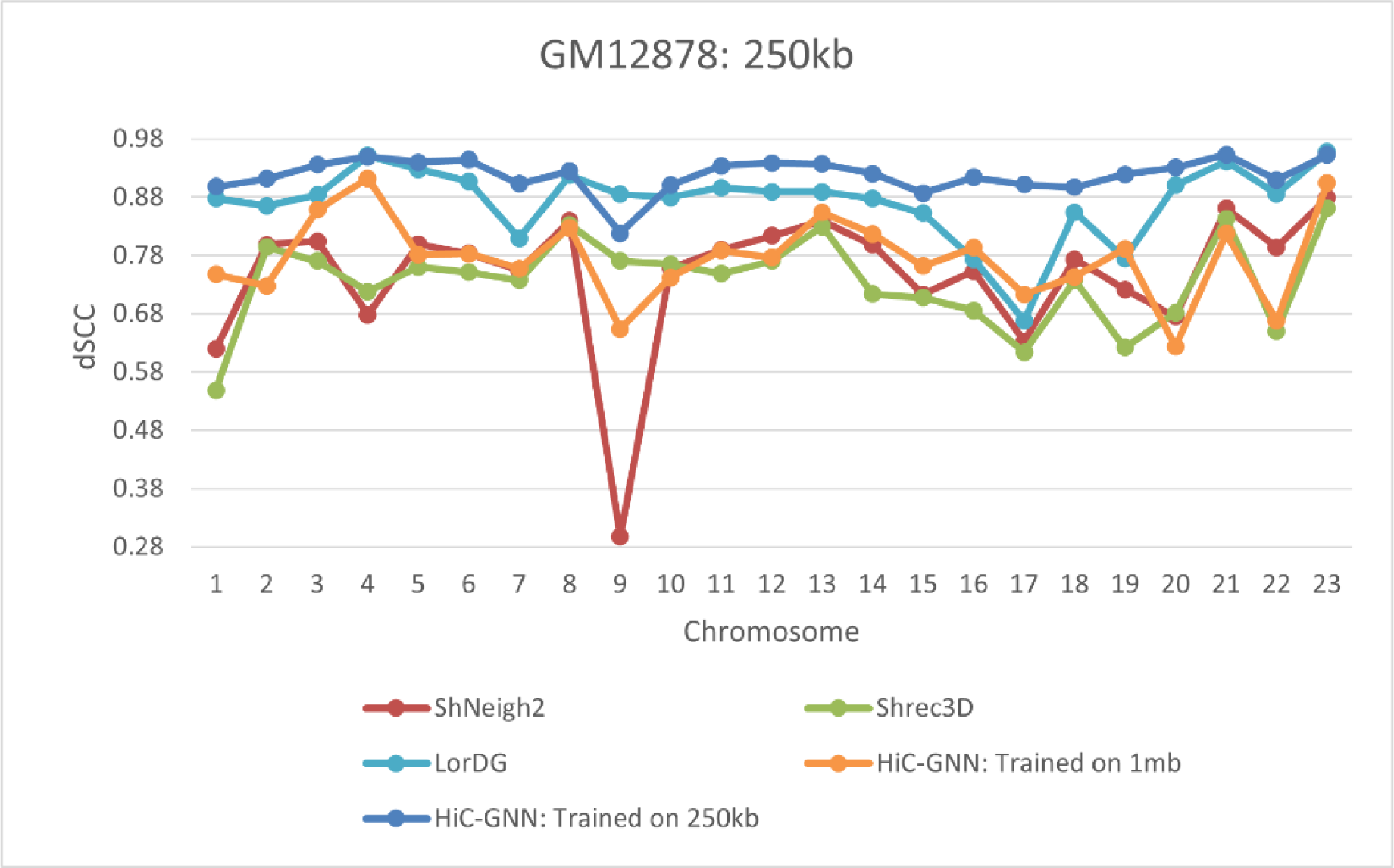
dSCC Comparison: 250kb Resolution. The figure shows a comparison of HiC-GNN with the other methods on the 250kb GM12878 cell data. The HiC-GNN models that were trained on the 250kb data are either on-par or outperform the other methods on the majority of the chromosomes. Although the HiC-GNN models that were trained on the 1mb data have lower performance, they still outperform ShNeigh and ShRec3D despite being trained on a quarter as many data instances.

**Figure 9.**
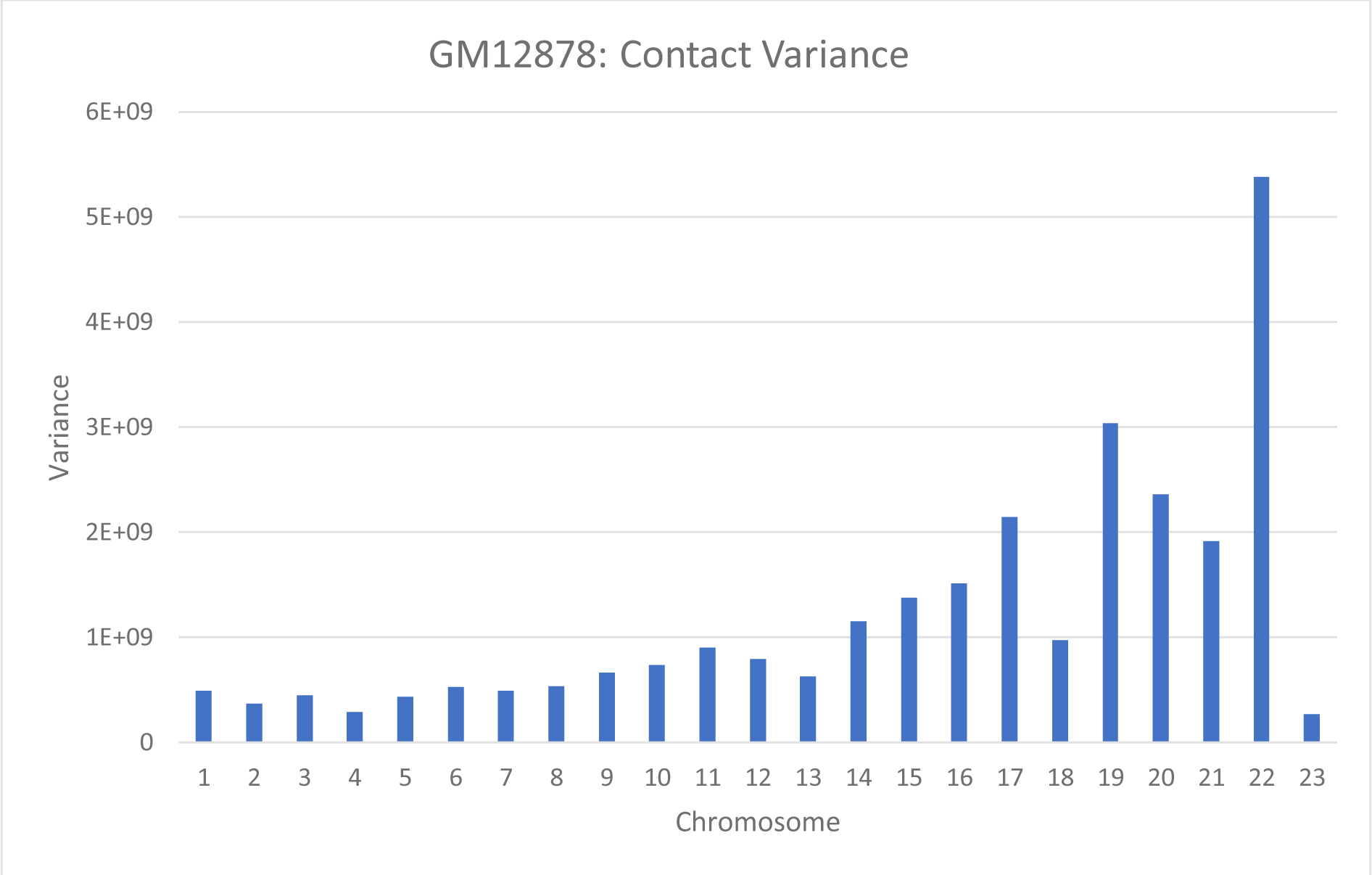
Contact Variances for GM12878 cell data at 1mb resolution. Chromosomes with higher contact variances lead to lower dSCC values for the other methods, whereas HiC-GNN is relatively robust to high contact variance.

**Fig 10.**
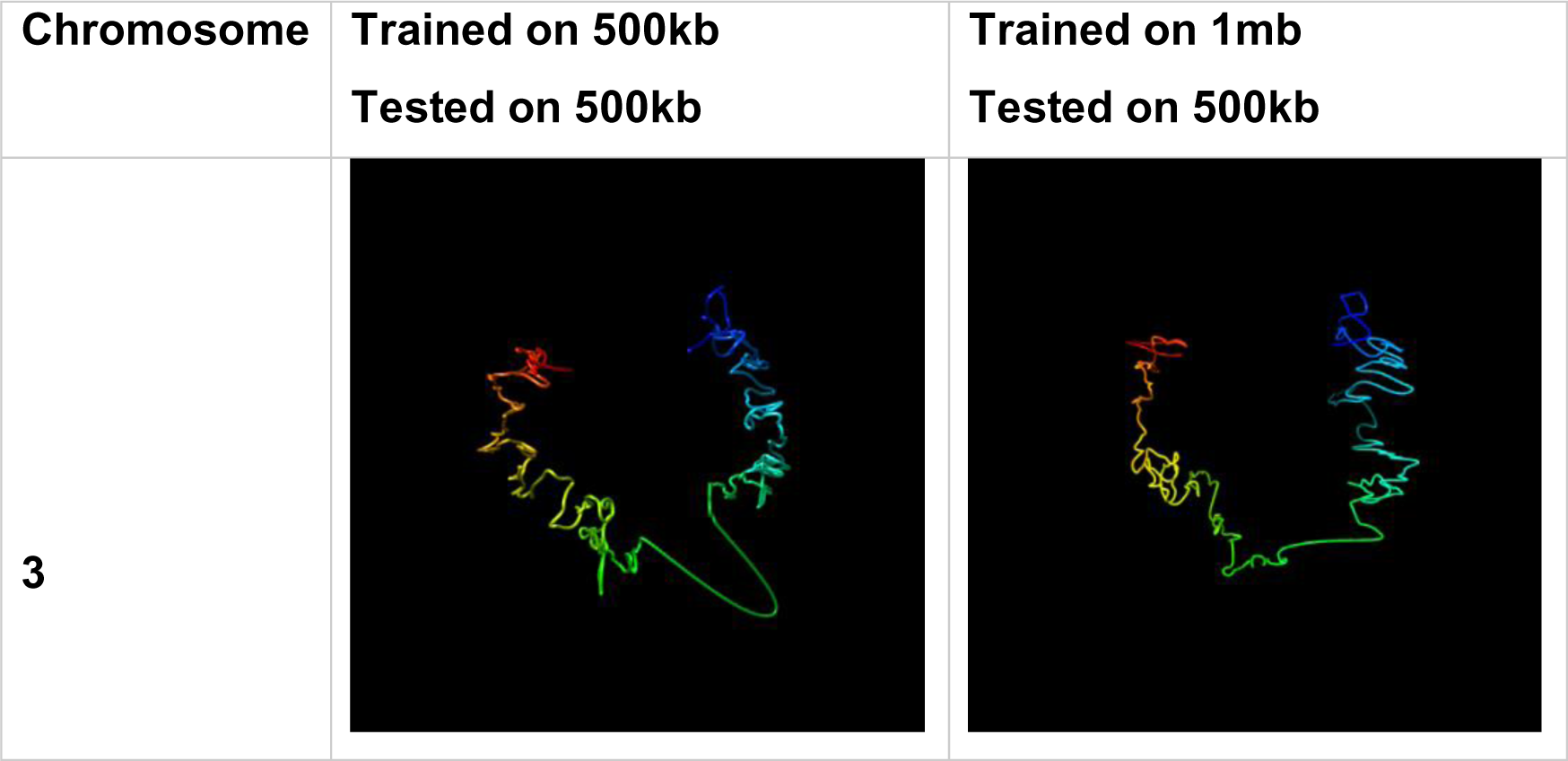
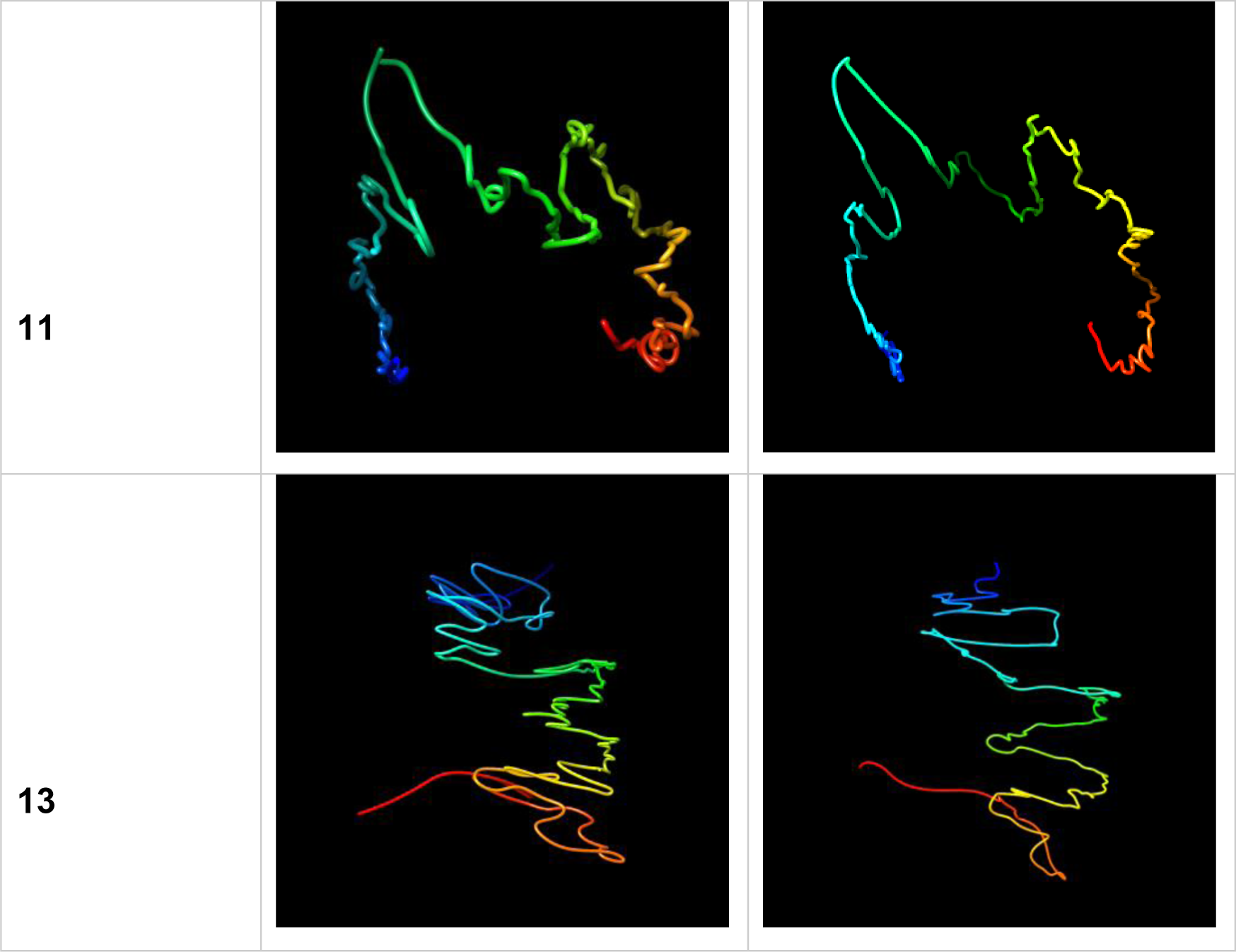
Visual Comparison of Structures Generated from HiC-GNN Generalized Across Resolution. The first column lists the chromosomes for which the 3D structure prediction was done, the second column shows the structures generated from a model trained and tested on a 500kb map and the third column shows structures generated from a model trained on a 1mb map and tested on a 500kb map.

**Fig 11.**
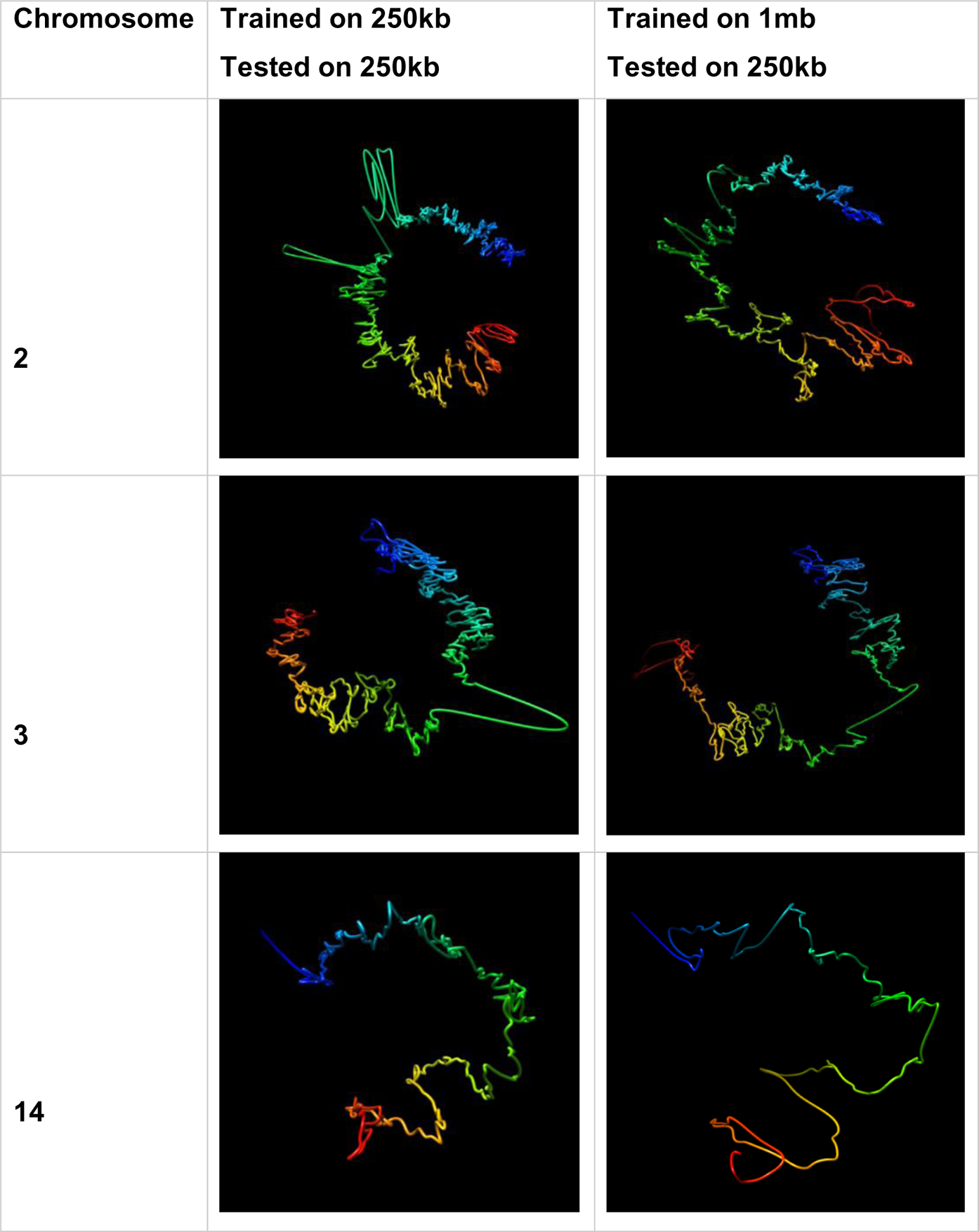
Visual Comparison of Structures Generated from HiC-GNN Generalized Across Resolution. The first column lists the chromosomes for which the 3D structure prediction was done, the second column shows the structures generated from a model trained and tested on a 250kb map and the third column shows structures generated from a model trained on a 1mb map and tested on a 250kb map.

#### Validation On ChIA-PET Data

Note that the results from Figs 6 through 8 imply a high correlation between the output model and the input wish distances. Thus, we know our method can accurately estimate 3D coordinates from a set of wish distances. Trussart et al.[27] showed that the dSCC between the distances corresponding to output models and the wish distances of the Hi-C map is a good proxy for model accuracy. Before discussing additional results involving dSCC, however, we further validate that our method does indeed produce representative models by comparing the results generated on the GM12878 cell line with orthogonal ChIA-PET data.

The ChIA-PET data provided by [37] consists of contact maps measuring interactions between the RNAPII complex in all 23 chromosomes of the GM12878 cell line. From these contact maps, RNAPII loops were identified by considering contact regions that have an interaction frequency greater than or equal to 5. To validate our method, we split this ChIA-PET data into two sets: one containing looped regions and one containing non-looped regions. We then calculated the distances of our output models between the identified looped and non-looped regions separately for each chromosome. If our models are representative of the true structure of the chromosome, then distances corresponding to looped regions of our output models should typically be smaller than distances corresponding to non-looped regions.

Figs 12 and 13 show the box plots for the looped and non-looped regions for all chromosomes combined for structures generated from both non-generalized and generalized HiC-GNN models at 500kb and 250kb resolution. From these figures, it is clear that the distribution of distances corresponding to the looped regions is centered around smaller values, thereby implying that the outputs of our method are consistent with the true structure of the chromosomes. This is true for both the generalized and non-generalized models.

**Fig 12.**
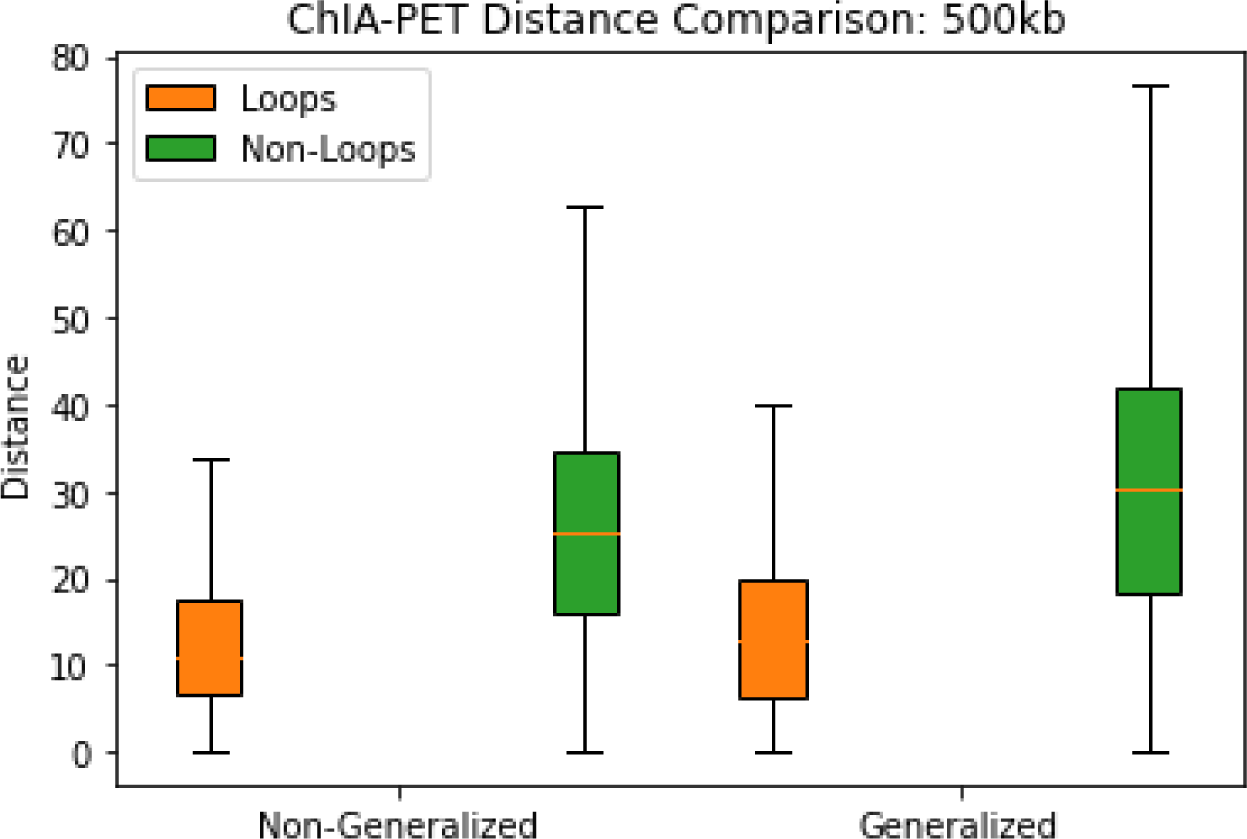
Result Validation on GM12879 500kb Resolution 3D Structure Predictions. The figure shows the box plots for the looped and non-looped regions for all chromosomes combined in the GM12879 cell line for generalized and non-generalized HiC-GNN models at 500kb resolution.

**Fig 13.**
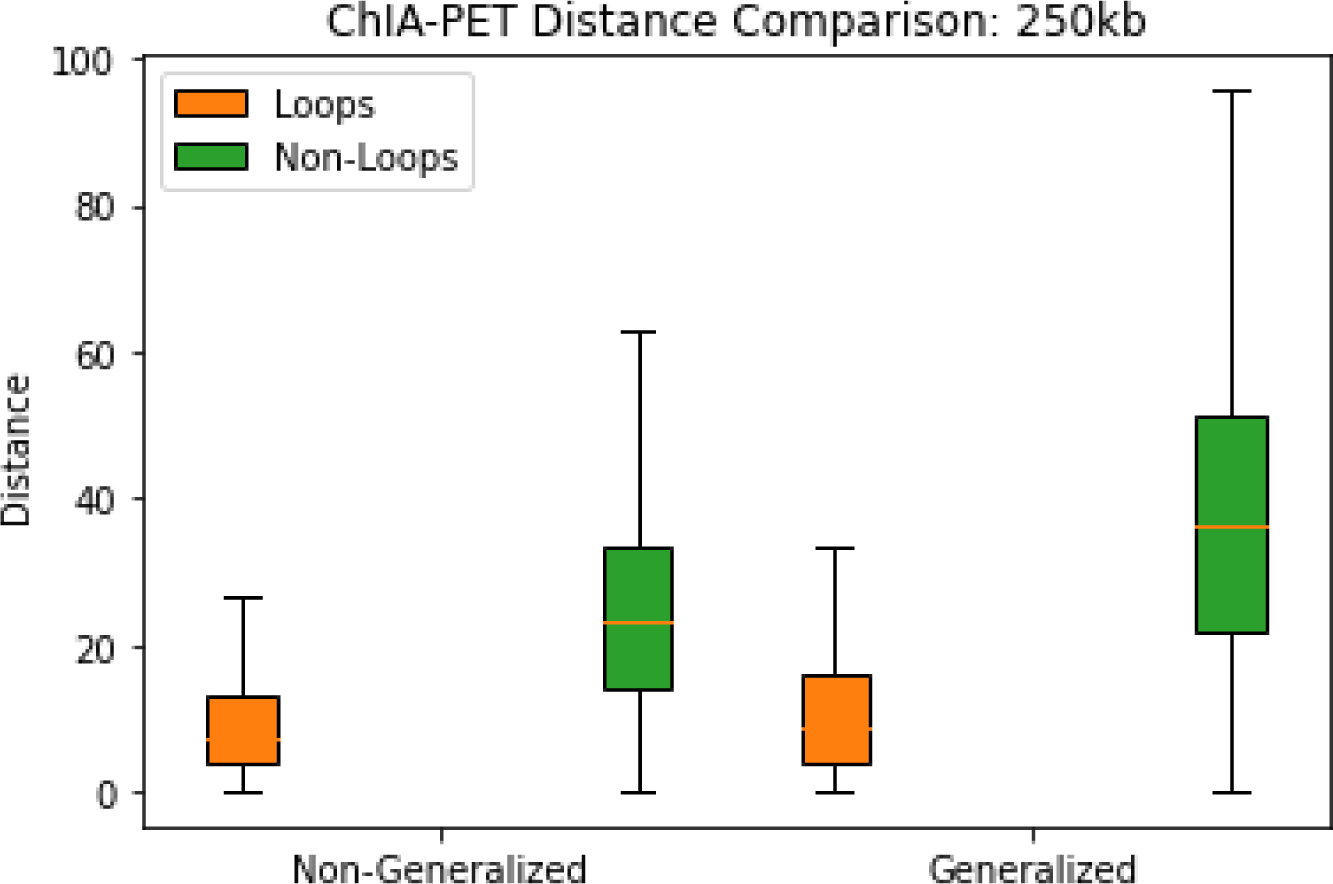
Result Validation on GM12879 250kb Resolution 3D Structure Predictions. The figure shows the box plots for the looped and non-looped regions for all chromosomes combined in the GM12879 cell line for generalized and non-generalized HiC-GNN models at 250kb resolution.

### GM06990

#### Generalization 2: Generalization Across Restriction Enzyme

To further test our method, we validated on the GM06990 cell line as well. This data consists of 22 Hi-C maps generated from the HindIII and Mbol restriction enzymes at 1mb resolution. We also tested how well HiC-GNN can generalize across input restriction enzymes. We tested this generalization by training a model on one restriction enzyme and testing on another, following the same alignment protocol as in the test for generalization across input resolution. The results of these tests along with comparisons with the other methods can be found in Fig 14 and 15.

**Fig 14.**
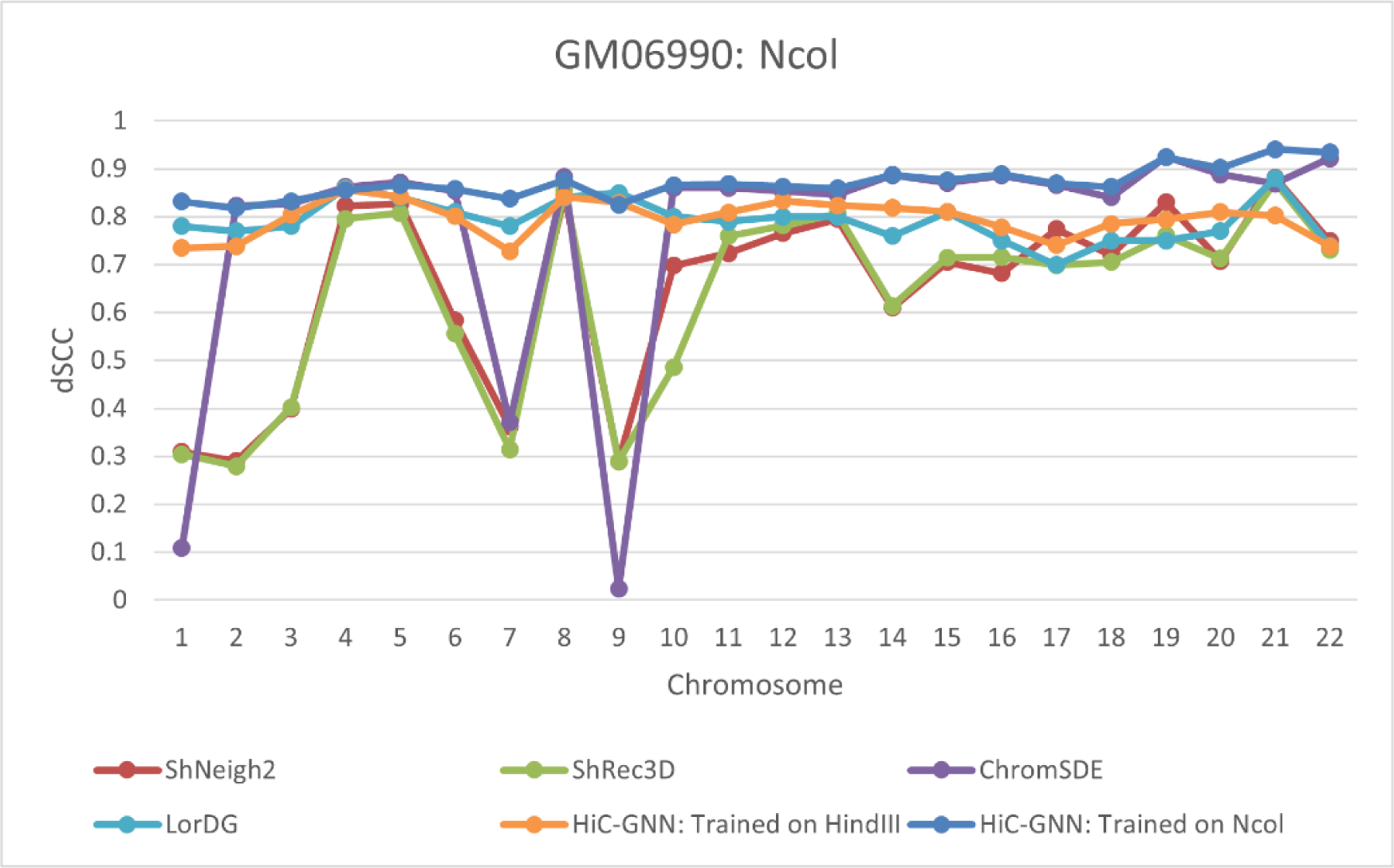
dSCC Comparison: Ncol Restriction Enzyme.

**Fig 15.**
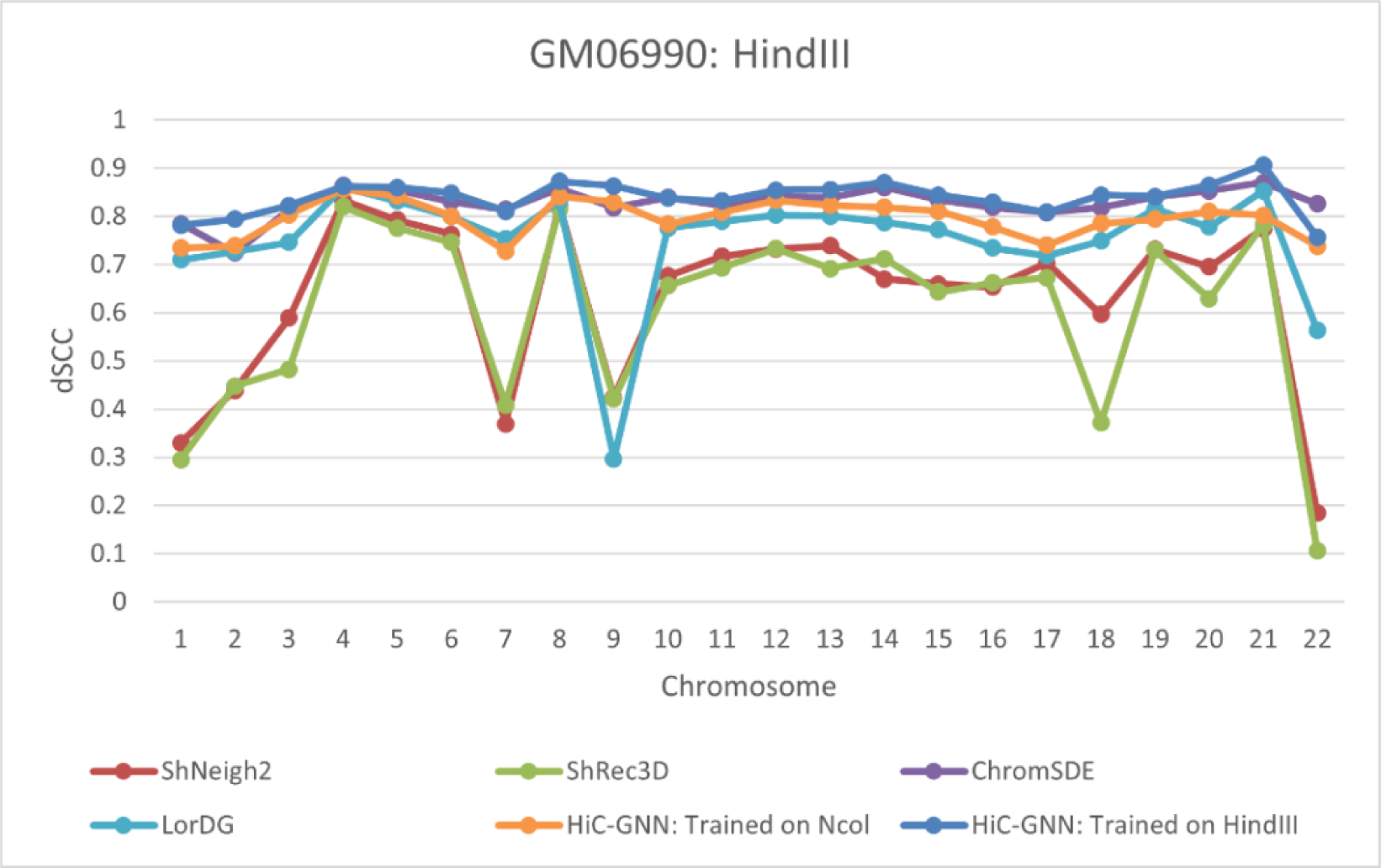
Comparison of the dSCC of output models for all three methods on GM0990 with the HindIII restriction enzyme.

From these figures, it is clear that the non-generalized HiC-GNN is either on par with or outperforms the other four methods. Moreover, the generalized HiC-GNN models outperform ShNeigh2, ShRec3D, and LorDG on most of the chromosomes despite being trained on data generated using an entirely different restriction enzyme. Figs 16 and 17 show the log variance for the contacts within these data sets. Once again, the performance of the other methods is severely affected by high variance in the input contact maps, particularly in chromosomes 9, 7, and 1 in the Ncol data and chromosomes 18, 9, and 1 in the HindIII data. Figs 18 and 19 provide a visual comparison between structures generated from a generalized HiC-GNN model and a non-generalized HiC-GNN model for three randomly selected chromosomes.

**Fig 16.**
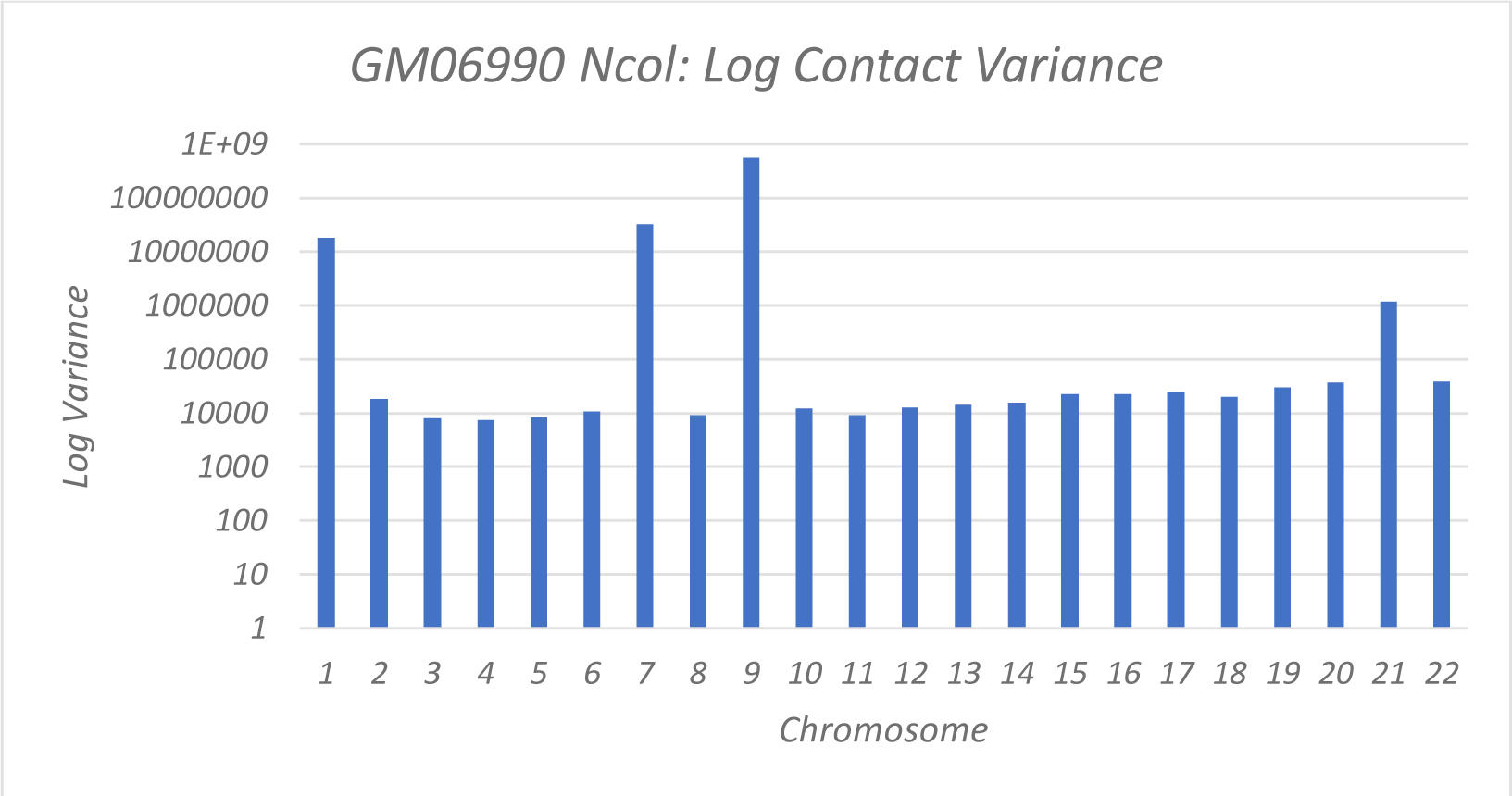
Log Contact Variances: GM06990 cell Ncol Data. Chromosomes with higher contact variances lead to lower dSCC values for the other methods, whereas HiC-GNN is relatively robust to high contact variance. This is particularly notable on chromosomes 1, 7, 9, and 21.

**Fig 17.**
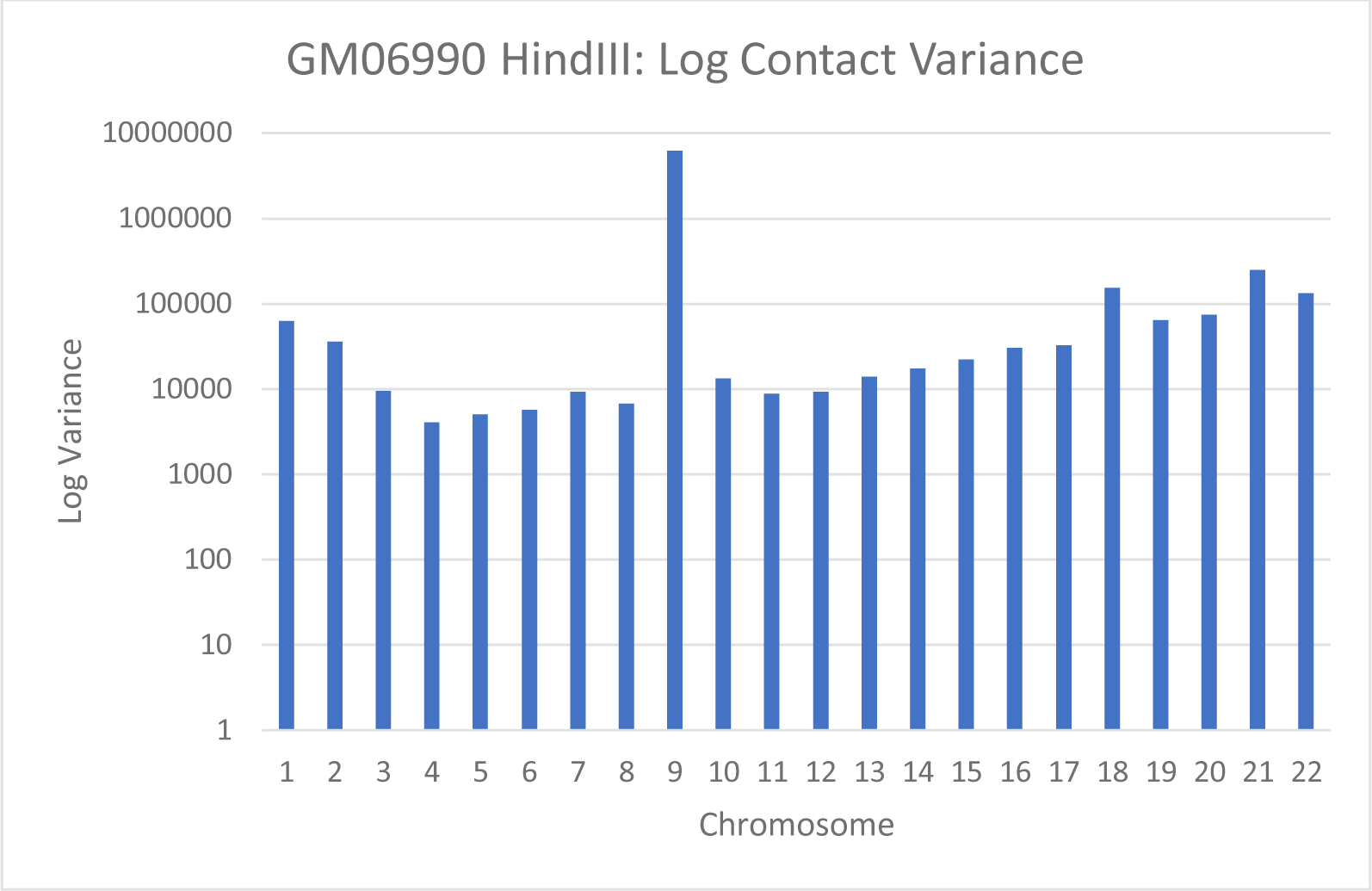
Log Contact Variances: GM06990 cell HindIII Data. Chromosomes with higher contact variances lead to lower dSCC values for the other methods, whereas HiC-GNN is relatively robust to high contact variance. This is particularly notable on chromosomes 1, 9 and 18.

**Fig 18.**
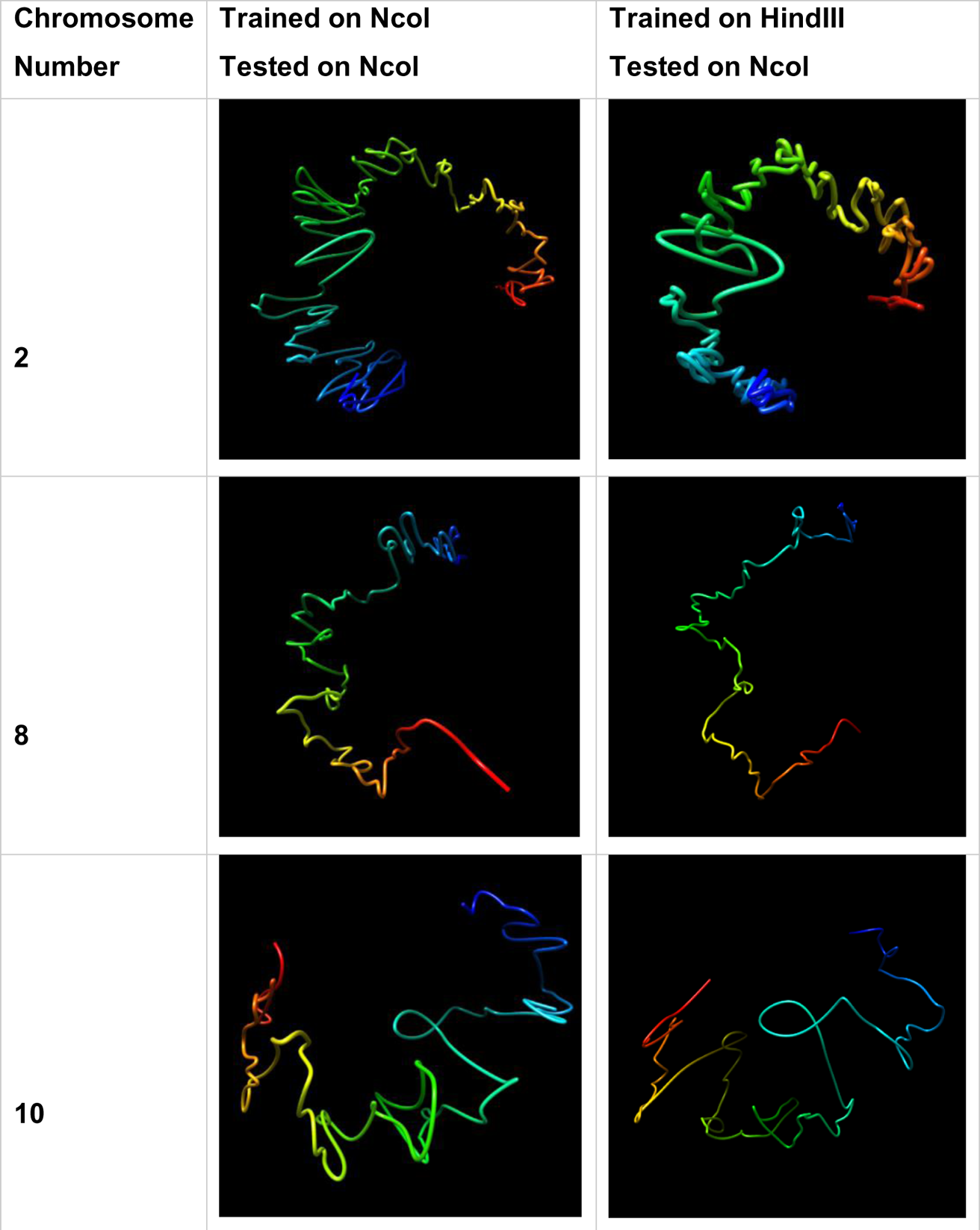
Visual Comparison of Structures Generated from HiC-GNN Generalized Across Resolution Enzymes. The first column lists the chromosomes for which the 3D structure prediction was done. The second column shows structures generated from a model trained and tested on the Ncol maps. The third column shows structures generated from a model trained on the HindIII maps and tested on the Ncol maps.

**Figure 19:**
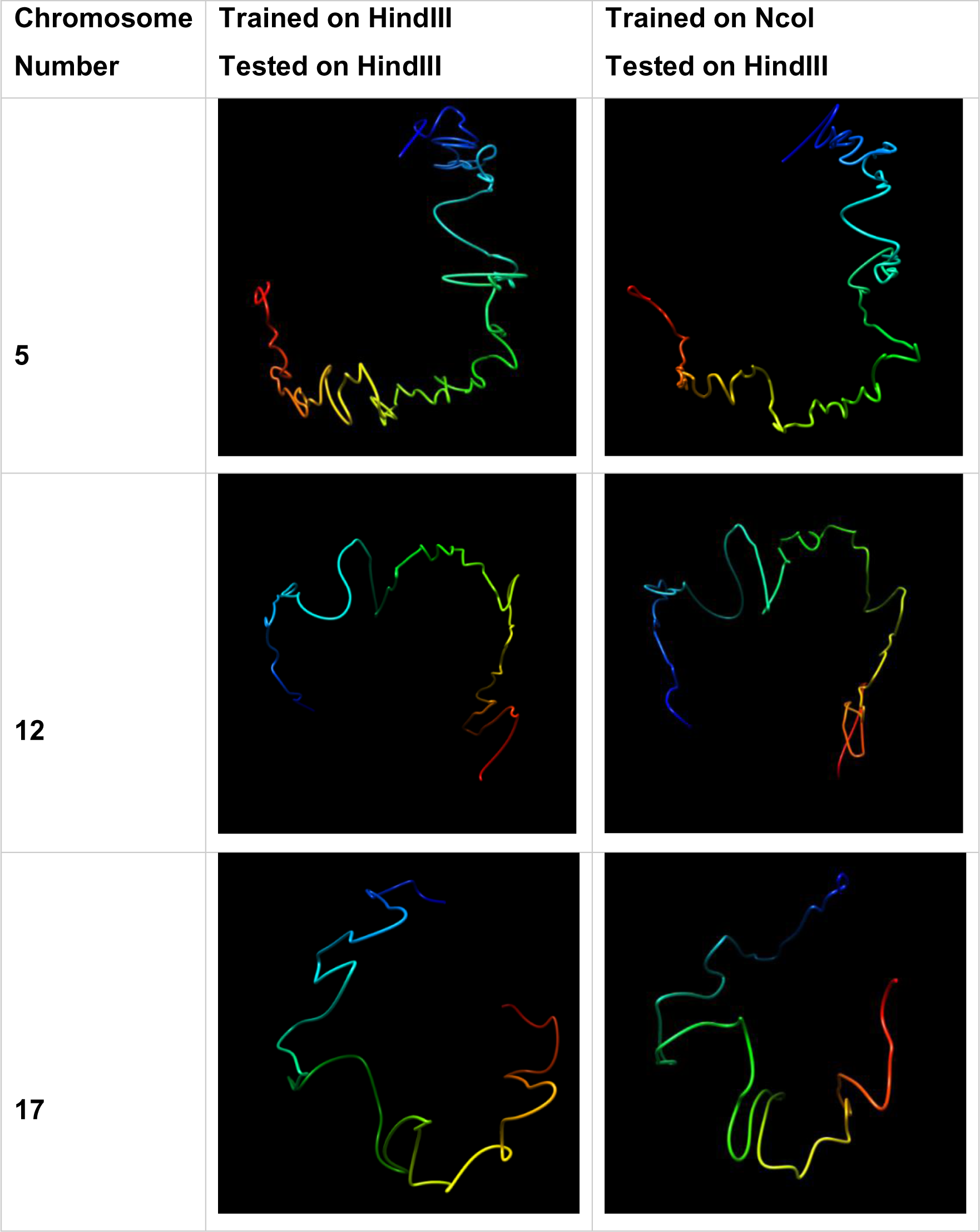
Visual Comparison of Structures Generated from HiC-GNN Generalized Across Resolution Enzymes. The first column lists the chromosomes for which the 3D structure prediction was done. The second column shows structures generated from a model trained and tested on the HindIII maps. The third column shows structures generated from a model trained on the Ncol maps and tested on the HindIII maps.

### K562

#### Generalization 3: Generalization Across Cell Populations

Finally, we tested how well HiC-GNN could generalize across different cell populations, i.e., how well can HiC-GNN perform when trained and tested across the Hi-C maps of chromosomes generated from separate Hi-C experiments. For this test, we utilized the K562 cell line from [32]. This data set consists of several sets of Hi-C maps generated from separate Hi-C experiments, each of which containing 23 chromosomes at 1mb resolution. We will refer to each of these orthogonal sets as *replicates*. Each replicate was generated using a separate population of cells. Due to variability in the size of the cell populations, each replicate has a varying quantity of total contacts across the entire genome. The smallest number of total contacts in the replicate sets is 53 million, and the largest number is 310 million. In most applications of Hi-C data analysis, it is typical to analyze the combination of all replicate maps corresponding to multiple Hi-C experiments. The combination of multiple maps simply refers to the matrix resulting from taking an element-wise sum of all replicates. By considering the combination of multiple replicate maps, the total amount of contacts to be analyzed is increased, thereby giving a more robust view of the distribution of the contact data.

We trained HiC-GNN on replicate maps and tested on the combined map for each chromosome within this data set to see how well HiC-GNN can generalize from contact spares data. To ensure a wide degree of data sparsity, we chose three replicate maps of varying numbers of total contacts. In addition to training models on these three replicates separately, we trained models on the map corresponding to their element-wise sum. The names of each replicate map along with their corresponding total contact number is given in Table 6. The results of this generalization can be found in Fig 20.

**Fig 20.**
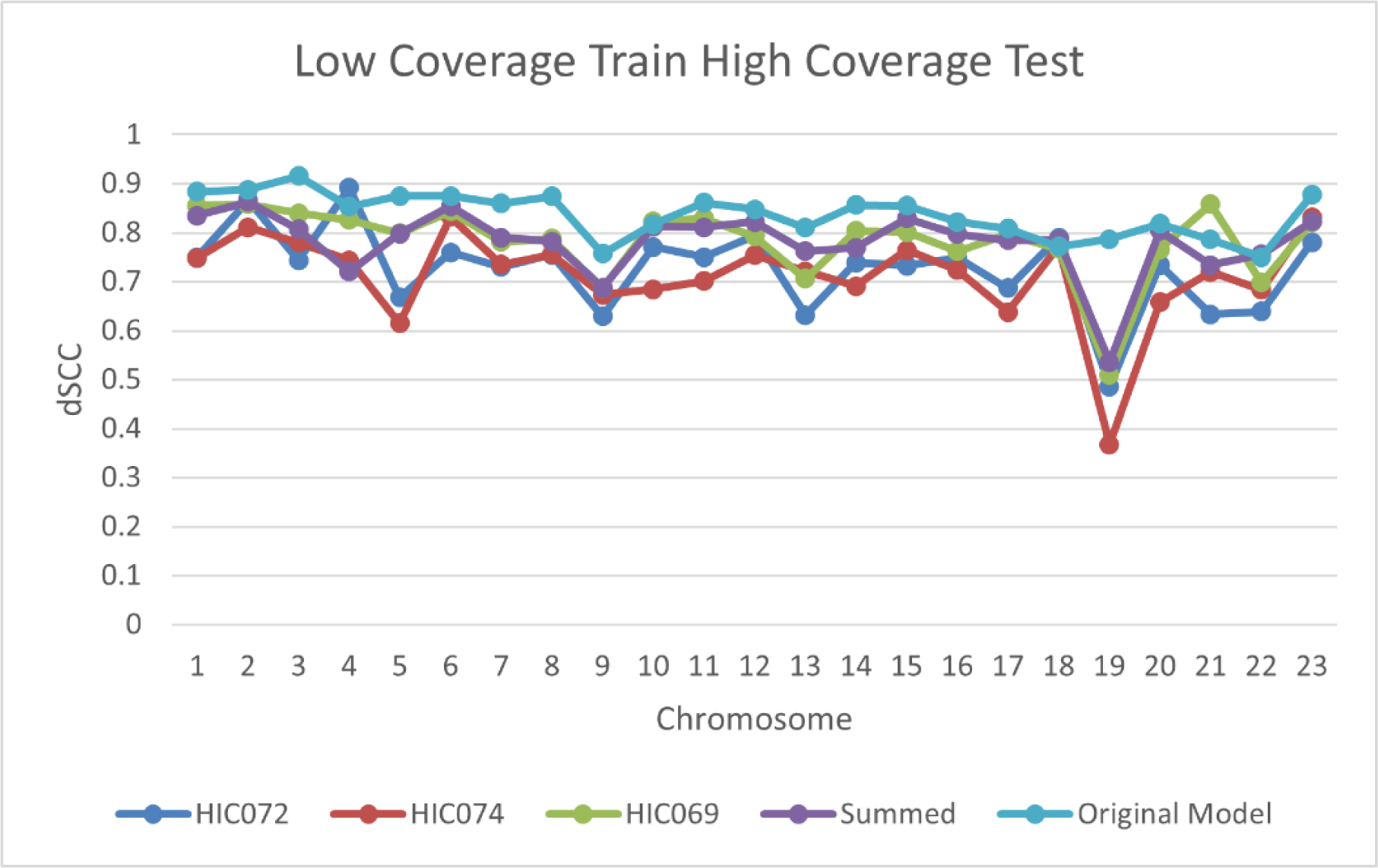
dSCC Comparison: Low to High Coverage. The figure shows the results of training HiC-GNN on maps with low coverage and testing on the combined map for each map in table. Despite the fact that the generalized models were trained on data containing as little as 6% of the contacts as the combined maps (in the case of HiC072), the reconstructive accuracy is still competitive with the original model.

**Table 6:** Names of each map and corresponding total number of contacts across the entire genome. The summed map corresponds to the element-wise sums of HiC02, HiC074, and HiC069. The combined map corresponds to the element-wise sums of each map within the dataset.

| Map Name | Number of Contacts (Millions) |
| --- | --- |
| HiC072 | 53 |
| HiC074 | 65 |
| HiC069 | 310 |
| Summed | 428 |
| Combined | 932 |

From these figures, one can tell that HiC-GNN can indeed generalize from maps containing higher degrees of contact sparsity. Although there is some drop in the reconstructive performance of HiC-GNN associated with generalizing on data containing fewer contacts, it is important to note that the generalized models were trained on data containing as little as less than 6% of the contacts (in the case of HiC072) as the combined maps. Fig 21 gives a visual comparison of the structures generated with generalized HiC-GNN models and structures generated using non-generalized HiC-GNN models (Trained on Summed and HiC072, Tested on Combined).

**Fig. 21:**
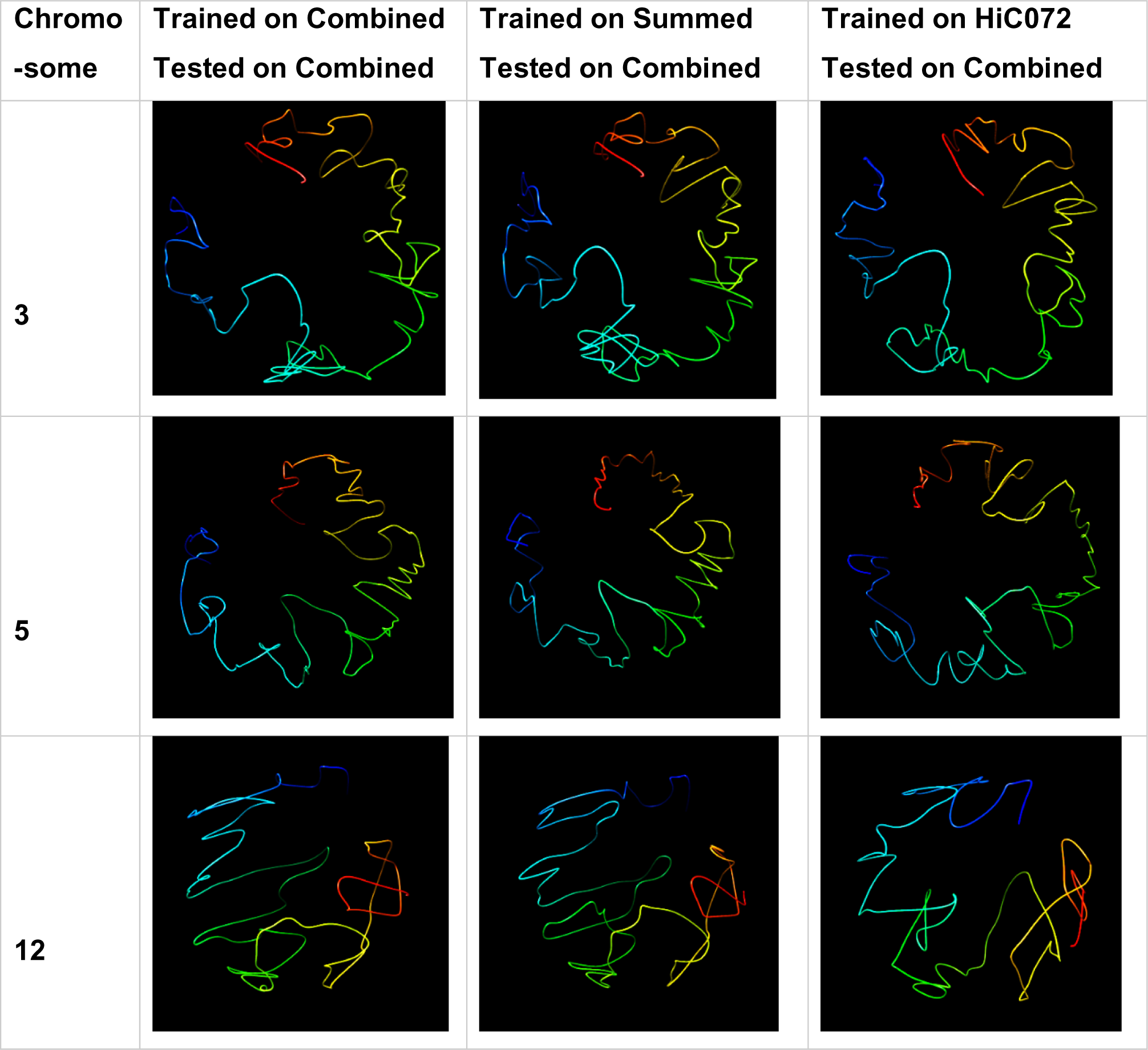
Visual Comparison of Structures Generated from HiC-GNN Generalized Across Cell Populations. The first column lists the chromosomes for which the 3D structure prediction was done, the second column shows the structures generated from a model trained and tested on a combined map, the third column shows structures generated from a model trained on a summed map and tested on a combined map, and the fourth column shows structures generated from a model trained on a HiC072 map and tested on a combined map.

## Discussion

In this paper, we presented a novel technique for predicting the 3D structure of chromosomes from Hi-C data using a node embedding algorithm and graph convolutional neural networks. Unlike other typical methods for chromosome structural inference, our method has the capability of generalizing across resolutions, restriction enzymes, and cell populations. We also showed that the performance of our method is superior when compared with other methods across multiple data sets. To our knowledge, the generalizations provided by our method are not present in any current methods for chromosome structure prediction.

The generalization of our method relies on an embedding alignment procedure. The increase in reconstructive accuracy following this embedding alignment procedure suggests that the node embeddings corresponding Hi-C maps are approximately isometric. This opens opportunities to use node embeddings as generalizable features for machine learning methods applied to other tasks involving Hi-C data. There are also several possible advantages to using GCNNs for the task of chromosome structure prediction that were not explored in this paper. For example, batching procedures could be used to improve the training process and more sophisticated embedding alignment could improve the reconstructive performance of generalized models.

## Supporting Information

S1 Table. Detailed list of configurations considered for hyperparameter tuning.

## Supporting information

S1 Table. Detailed list of configurations considered for hyperparameter tuning.

## References

1. T. Misteli, “Beyond the sequence: cellular organization of genome function,” Cell, vol. 128, p. 787–800, 2007.

2. P. Fraser and W. Bickmore, “Nuclear organization of the genome and the potential for gene regulation,” Nature, vol. 447, p. 413–417, 2007.

3. J. Dekker, “Gene regulation in the third dimension,” Science, vol. 319, p. 1793–1794, 2008.

4. J. Dekker, K. Rippe, M. Dekker and N. Kleckner, “Capturing chromosome conformation,” Science, vol. 295, p. 1306–1311, 2002.

5. M. Simonis, P. Klous, E. Splinter, Y. Moshkin, R. Willemsen, E. De Wit, B. Van Steensel and W. De Laat, “Nuclear organization of active and inactive chromatin domains uncovered by chromosome conformation capture–on-chip (4C),” Nature Genetics, vol. 38, p. 1348–1354, 2006.

6. J. Dostie, T. A. Richmond, R. A. Arnaout, R. R. Selzer, W. L. Lee, T. A. Honan, E. D. Rubio, A. Krumm, J. Lamb, C. Nusbaum and others, “Chromosome Conformation Capture Carbon Copy (5C): a massively parallel solution for mapping interactions between genomic elements,” Genome Research, vol. 16, p. 1299–1309, 2006.

7. N. L. Van Berkum, E. Lieberman-Aiden, L. Williams, M. Imakaev, A. Gnirke, L. A. Mirny, J. Dekker and E. S. Lander, “Hi-C: a method to study the three-dimensional architecture of genomes.,” JoVE (Journal of Visualized Experiments), p. e1869, 2010.

8. E. De Wit and W. De Laat, “A decade of 3C technologies: insights into nuclear organization,” Genes & Development, vol. 26, p. 11–24, 2012.

9. E. Lieberman-Aiden, N. L. Van Berkum, L. Williams, M. Imakaev, T. Ragoczy, A. Telling, I. Amit, B. R. Lajoie, P. J. Sabo, M. O. Dorschner and others, “Comprehensive mapping of long-range interactions reveals folding principles of the human genome,” Science, vol. 326, p. 289–293, 2009.

10. O. Oluwadare, M. Highsmith and J. Cheng, “An overview of methods for reconstructing 3-D chromosome and genome structures from Hi-C data,” Biological Procedures Online, vol. 21, p. 1–20, 2019.

11. T. Trieu and J. Cheng, “3D genome structure modeling by Lorentzian objective function,” Nucleic Acids Research, vol. 45, p. 1049–1058, 2017.

12. O. Oluwadare, Y. Zhang and J. Cheng, “A maximum likelihood algorithm for reconstructing 3D structures of human chromosomes from chromosomal contact data,” BMC Genomics, vol. 19, p. 1–17, 2018.

13. L. Rieber and S. Mahony, “miniMDS: 3D structural inference from high-resolution Hi-C data,” Bioinformatics, vol. 33, p. i261–i266, 2017.

14. T. Trieu, O. Oluwadare and J. Cheng, “Hierarchical reconstruction of high-resolution 3D models of large chromosomes,” Scientific Reports, vol. 9, p. 1–12, 2019.

15. B. Adhikari, T. Trieu and J. Cheng, “Chromosome3D: reconstructing three-dimensional chromosomal structures from Hi-C interaction frequency data using distance geometry simulated annealing,” BMC Genomics, vol. 17, p. 1–9, 2016.

16. Z. Zhang, G. Li, K.-C. Toh and W.-K. Sung, “Inference of spatial organizations of chromosomes using semi-definite embedding approach and Hi-C data,” in Annual international conference on research in computational molecular biology, 2013.

17. S. Sazer and H. Schiessel, “The biology and polymer physics underlying large-scale chromosome organization,” Traffic, vol. 19, p. 87–104, 2018.

18. A. Lesne, J. Riposo, P. Roger, A. Cournac and J. Mozziconacci, “3D genome reconstruction from chromosomal contacts,” Nature Methods, vol. 11, p. 1141–1143, 2014.

19. F.-Z. Li, Z.-E. Liu, X.-Y. Li, L.-M. Bu, H.-X. Bu, H. Liu and C.-M. Zhang, “Chromatin 3D structure reconstruction with consideration of adjacency relationship among genomic loci,” BMC Bioinformatics, vol. 21, p. 1–17, 2020.

20. P. A. Knight and D. Ruiz, “A fast algorithm for matrix balancing,” IMA Journal of Numerical Analysis, vol. 33, p. 1029–1047, 2013.

21. H. Lyu, E. Liu and Z. Wu, “Comparison of normalization methods for Hi-C data,” BioTechniques, vol. 68, p. 56–64, 2020.

22. D. P. Kingma and J. Ba, “Adam: A method for stochastic optimization,” *arXiv preprint arXiv:1412*.6980, 2014

23. A. Pombo and M. Nicodemi, “Physical mechanisms behind the large scale features of chromatin organization,” Transcription, vol. 5, p. e28447, 2014.

24. M. Barbieri, M. Chotalia, J. Fraser, L.-M. Lavitas, J. Dostie, A. Pombo and M. Nicodemi, “Complexity of chromatin folding is captured by the strings and binders switch model,” Proceedings of the National Academy of Sciences, vol. 109, p. 16173–16178, 2012.

25. A. M. Chiariello, C. Annunziatella, S. Bianco, A. Esposito and M. Nicodemi, “Polymer physics of chromosome large-scale 3D organisation,” Scientific Reports, vol. 6, p. 1–8, 2016.

26. J. Mateos-Langerak, M. Bohn, W. de Leeuw, O. Giromus, E. M. M. Manders, P. J. Verschure, M. H. G. Indemans, H. J. Gierman, D. W. Heermann, R. Van Driel and others, “Spatially confined folding of chromatin in the interphase nucleus,” Proceedings of the National Academy of Sciences, vol. 106, p. 3812–3817, 2009.

27. M. Trussart, F. Serra, D. Bau, I. Junier, L. Serrano and M. A. Marti-Renom, “Assessing the limits of restraint-based 3D modeling of genomes and genomic domains,” Nucleic Acids Research, vol. 43, p. 3465–3477, 2015.

28. J. Tang, M. Qu, M. Wang, M. Zhang, J. Yan and Q. Mei, “Line: Large-scale information network embedding,” in Proceedings of the 24th international conference on world wide web, 2015.

29. H. Ashoor, X. Chen, W. Rosikiewicz, J. Wang, A. Cheng, P. Wang, Y. Ruan and S. Li, “Graph embedding and unsupervised learning predict genomic sub-compartments from HiC chromatin interaction data,” Nature Communications, vol. 11, p. 1–11, 2020.

30. J. C. Gower, G. B. Dijksterhuis and others, Procrustes problems, vol. 30, Oxford University Press on Demand, 2004.

31. P. H. Schönemann, “A generalized solution of the orthogonal procrustes problem,” Psychometrika, vol. 31, p. 1–10, 1966.

32. S. S. P. Rao, M. H. Huntley, N. C. Durand, E. K. Stamenova, I. D. Bochkov, J. T. Robinson, A. Sanborn, I. Machol, A. D. Omer, E. S. Lander and E. L. Aiden, “A three-dimensional map of the human genome at kilobase resolution reveals principles of chromatin looping,” Cell (Cambridge), vol. 159, pp. 1665–1680, 2014.

33. O. Oluwadare, M. Highsmith, D. Turner, E. Lieberman Aiden and J. Cheng, “GSDB: a database of 3D chromosome and genome structures reconstructed from Hi-C data,” BMC Molecular and Cell Biology, vol. 21, pp. 60–60, 2020.

34. N. C. Durand, J. T. Robinson, M. S. Shamim, I. Machol, J. P. Mesirov, E. S. Lander and E. L. Aiden, “Juicebox Provides a Visualization System for Hi-C Contact Maps with Unlimited Zoom,” Cell Systems, vol. 3, pp. 99–101, 2016.

35. M. F. Carey, C. L. Peterson and S. T. Smale, “Chromatin immunoprecipitation (chip),” Cold Spring Harbor Protocols, vol. 2009, p. pdb–prot5279, 2009.

36. G. Li, L. Cai, H. Chang, P. Hong, Q. Zhou, E. V. Kulakova, N. A. Kolchanov and Y. Ruan, “Chromatin Interaction Analysis with Paired-End Tag (ChIA-PET) sequencing technology and application,” BMC Genomics, vol. 15 Suppl 12, pp. S11–S11, 2014.

37. T. Barrett, T. O. Suzek, D. B. Troup, S. E. Wilhite, W.-C. Ngau, P. Ledoux, D. Rudnev, A. E. Lash, W. Fujibuchi and R. Edgar, “NCBI GEO: mining millions of expression profilesâ€”database and tools,” Nucleic Acids Research, vol. 33, pp. D562–D566, 2005;2004;.

38. K. Xu, W. Hu, J. Leskovec and S. Jegelka, “How powerful are graph neural networks?,” arXiv preprint arXiv:1810.00826, 2018.

39. Y. Wang, Y. Sun, Z. Liu, S. E. Sarma, M. M. Bronstein and J. M. Solomon, “Dynamic graph cnn for learning on point clouds,” Acm Transactions On Graphics (TOG), vol. 38, p. 1–12, 2019.

40. M. Rousseau, J. Fraser, M. A. Ferraiuolo, J. Dostie and M. Blanchette, “Three-dimensional modeling of chromatin structure from interaction frequency data using Markov chain Monte Carlo sampling,” BMC Bioinformatics, vol. 12, p. 1–16, 2011.

41. Y. Li, D. Tarlow, M. Brockschmidt and R. Zemel, “Gated graph sequence neural networks,” arXiv preprint arXiv:1511.05493, 2015.

42. T. N. Kipf and M. Welling, “Semi-supervised classification with graph convolutional networks,” arXiv preprint arXiv:1609.02907, 2016.

43. W. L. Hamilton, R. Ying and J. Leskovec, “Inductive representation learning on large graphs,” in Proceedings of the 31st International Conference on Neural Information Processing Systems, 2017.

44. B. Fernando, A. Habrard, M. Sebban and T. Tuytelaars, “Unsupervised visual domain adaptation using subspace alignment,” in Proceedings of the IEEE international conference on computer vision, 2013.

45. J. Du, S. Zhang, G. Wu, J. M. F. Moura and S. Kar, “Topology adaptive graph convolutional networks,” arXiv preprint arXiv:1710.10370, 2017.

